# Value Certainty in Drift-Diffusion Models of Preferential Choice

**DOI:** 10.1101/2020.08.22.262725

**Authors:** Douglas Lee, Marius Usher

## Abstract

The *drift-diffusion model* (DDM) is widely used and broadly accepted for its ability to account for binary choices (in both the perceptual and preferential domains) and response times (RT), as a function of the stimulus or the choice alternative (or option) values. The DDM is built on an evidence accumulation-to-bound concept, where, in the value domain, a decision maker repeatedly samples the mental representations of the values of the available options until satisfied that there is enough evidence (or support) in favor of one option over the other. As the signals that drive the evidence are derived from value estimates that are not known with certainty, repeated sequential samples are necessary to average out noise. The classic DDM does not allow for different options to have different levels of precision in their value representations. However, recent studies have shown that decision makers often report levels of certainty regarding value estimates that vary across choice options. There is therefore a need to extend the DDM to include an option-specific value certainty component. We present several such DDM extensions and validate them against empirical data from four previous studies. The data support best a DDM version in which the drift of the accumulation is based on a sort of signal-to-noise ratio of value for each option (rather than a mere accumulation of samples from the corresponding value distributions). This DDM variant accounts for the impact of value certainty on both choice consistency and response time present in the empirical data.

## Main

The *drift-diffusion model* (DDM; Bogacz et al, 2006; Glickman & Usher, 2019; Gold & Shadlen, 2007; Ratcliff, 1978; Ratcliff & Rouder, 1998; Ratcliff & McKoon, 2008; Stine et al, 2020) is ubiquitous in the contemporary literature on decision making, including research spanning the fields of psychology, neuroscience, economics, and consumer behavior. The DDM is a parsimonious mechanism that explains normative decisions by averaging out noise in value comparisons (e.g., stochastic fluctuations in value signals that may result from variability in the firing of neural populations, from variability in attention or in memory retrieval of the values associated with the choice alternatives, or from variability intrinsic in the environment when the alternatives are characterized by fluctuating or stochastic values; e.g., Hertwig, Barron, & Weber, 2004; Glickman & Usher, 2019). This DDM mechanism implements an optimal stopping rule (optimizing response time (RT) for a specified accuracy; Wald, 1948; Gold & Shadlen, 2007; Bogacz et al, 2006). Moreover, this model accounts well for the dependency of the RT distribution on the values (or stimulus strength) of the choice options (Ratcliff & Rouder, 1998; Ratcliff & McKoon, 2008), and for the *speed-accuracy* tradeoff — the often-observed phenomenon that decision makers are able to improve their choice accuracy by taking longer to decide (on average; Wickelgren, 1977) — which is explained by an expansion of the DDM response boundaries.

While initially used in the domain of perceptual (Ratcliff & Rouder, 1998) or memory-based decisions (Ratcliff, 1978), the DDM core principles of sequential sampling and integration to boundary have since become a central component of models of preference-based decisions (Busemeyer & Townsend, 1993; Busemeyer & Diederich, 2002; Busemeyer et al, 2019; Fudenberg, Strack, & Strzalecki, 2018; Tajima, Drugowitsch, & Pouget, 2016; Polania et al, 2015; Philiastides & Ratcliff, 2013; Milosavljevic et al, 2010; Krajbich, Armel, & Rangel, 2010; Basten et al, 2010; Roe, Busemeyer & Townsend, 2001; Turner, Schley, Muller, & Tsetsos, 2018; Usher & McClelland, 2004). For example, in the *Decision Field Theory* (DFT) model (Busemeyer & Townsend, 1993; Busemeyer & Diederich, 2002; Busemeyer et al, 2019; Roe et al., 2001) — one of the first models that applied sequential sampling principles to preferential choice — it is assumed that variability in the value integration corresponds to fluctuations of attention between the relevant attributes (or outcomes) that characterize the alternatives. In a more recent application of the DDM to preferential choice, the drift rate is assumed to be modulated by visual attention across the alternatives (accordingly, this model was labeled the *attentional drift-diffusion model* or aDDM), which explains why people tend to choose more often the alternatives that they look at longer (Krajbich, Armel, & Rangel, 2010). In two more recent studies (Tajima et al; 2016; Fudenberg, Strack, & Strzalecki, 2018), the authors carried out normative analyses of the optimal decision policy in preferential choice. Both of these studies were based on principles of Bayesian updating of value estimates and optimal stopping policies, given the cost of accumulating evidence. Tajima et al (2016) converged on the conclusion that “similar to perceptual decisions, drift diffusion models implement the optimal strategy for value-based decisions.” An important assumption for the derivation of this optimal policy is that “the value of each option is represented by a probability distribution whose mean is the true (subjective) value of the option, and whose variance corresponds to sampling noise or uncertainty about the true value.” This noise could be interpreted as imprecision in the value representations themselves. Similarly, the model of Fudenberg et al (2018) includes a common variance term meant to represent the uncertainty about the option value estimates (prior to accumulating evidence, which is reduced as more evidence is accumulated). Thus, the momentary signal about the relative value of the options (the so-called evidence for one option over the other) fluctuates randomly around a fixed value (the so-called drift rate, which corresponds to the difference in the means of the two value distributions). In order to average out this noise, the values of the alternatives are repeatedly sampled (possibly from memory; Shadlen & Shohamy, 2016; Bakkour et al, 2019) and accumulated over time until a sufficient amount of evidence has been acquired to allow for a choice to be made. Accordingly, the DDM includes response boundaries for each option (typically symmetric) that trigger a choice once reached by the evidence accumulator. Thus, the fundamental components of the drift-diffusion process are the drift rate (proportional to the difference in the option values), the diffusion coefficient (the degree of stochasticity of the system), and the choice boundaries (the threshold of minimum required evidence for a given level of caution, which controls the speed-accuracy tradeoff; see Figure 1).^1^

**Figure 1:**
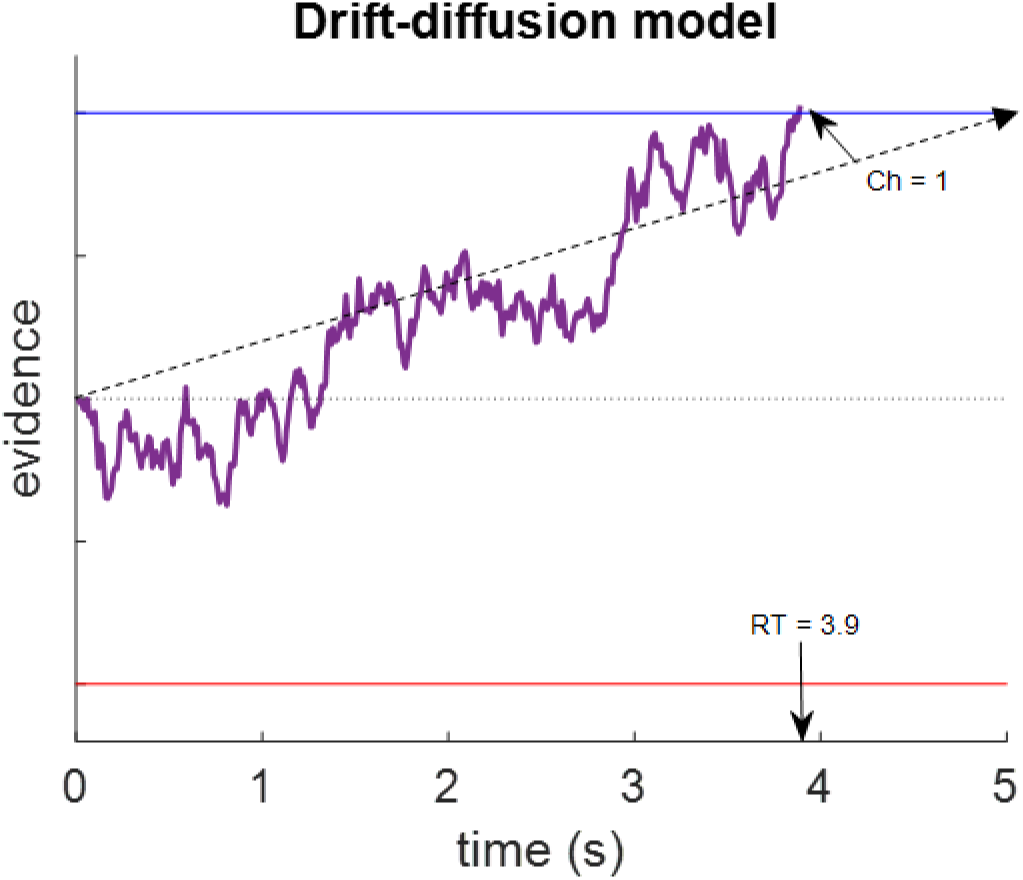
An illustration of the classic DDM. Evidence accumulates across time, following a fixed drift trajectory (black dashed arrow, whose slope corresponds to the value difference of the choice options multiplied by a scalar coefficient), corrupted by white processing or sampling noise. Here, the accumulated evidence reaches the upper boundary after 3.9 seconds, and a choice for option 1 is recorded.

The classic DDM implicitly assumes that processing (or sampling) noise is independent of the identity of the options contained in a particular choice set. That is to say, the sampling noise in the DDM is not option-specific; most of the DDM applications to preference-based decisions have assumed no option-specific noise (but see Ratcliff, Voskuilen, & Teodorescu, 2018; Teodorescu, Moran, & Usher, 2016 for DDM variants in which the noise increases with task difficulty; and Ratcliff & McKoon, 2018, for an application of option-specific variability in decisions about numerosity). However, the brain is known to encode not only the subjective value of options, but also the subjective certainty about the value (Lebreton et al, 2015). Efficient coding principles also suggest that the values of options will be represented with greater precision when those options are encountered more frequently (Heng, Woodford, & Polania, 2020). It is thus reasonable to suggest that the representations of value that the brain uses to inform the decision process vary (Tajima et al, 2016), and that the degree of imprecision (or uncertainty) is not the same for all choice options. Indeed, it has been shown that decision makers hold highly variable beliefs about the certainty of their value estimates for different options, and that those beliefs are relatively stable within individuals (Lee & Daunizeau, 2020, 2021; Lee & Coricelli, 2020; Gwinn & Krajbich, 2020; Polania, Woodford, & Ruff, 2019). It has further been shown that the variability in value (un)certainty has a clear and systematic impact on both choice and RT (Lee & Daunizeau, 2020, 2021; Lee & Coricelli, 2020). Specifically, the judged value certainty, C, correlates positively with choice consistency (the equivalent of choice accuracy for preferential choice) and negatively with RT (see Figure 2) ^2^ . Furthermore, the difference in certainty (in favor of the higher-valued alternative), dC, is also associated with higher choice consistency and faster RT (see Figure 2). This provides a qualitative benchmark that any DDM variant that includes option-specific certainty should be able to account for. To summarize these previous findings, we pooled the data from four studies (n=191) in which participants made choices between food items after having had rated their subjective value of the items and their certainty about their ratings. We then conducted mixed model regression analyses: logistic for consistency, linear for log(RT). The random effects variables were the different studies and individual participants, and the fixed effects variables were dV, C, and dC (see Figure 2). We focused here on the effects of dV, C, and dC — the former is expected in any DDM, while the latter two are challenges that we aim to account for. The actual data also shows an effect of the overall value (V = V1 + V2) on RT (see Figure S5 in the Supplementary Material). Such dependence is usually not obtained in DDM type models, and since our focus here is on effects of value certainty, we defer this to the Discussion section; as shown in the Supplementary Material, the DDM extension that we propose does account for this additional overall value effect.

**Figure 2:**
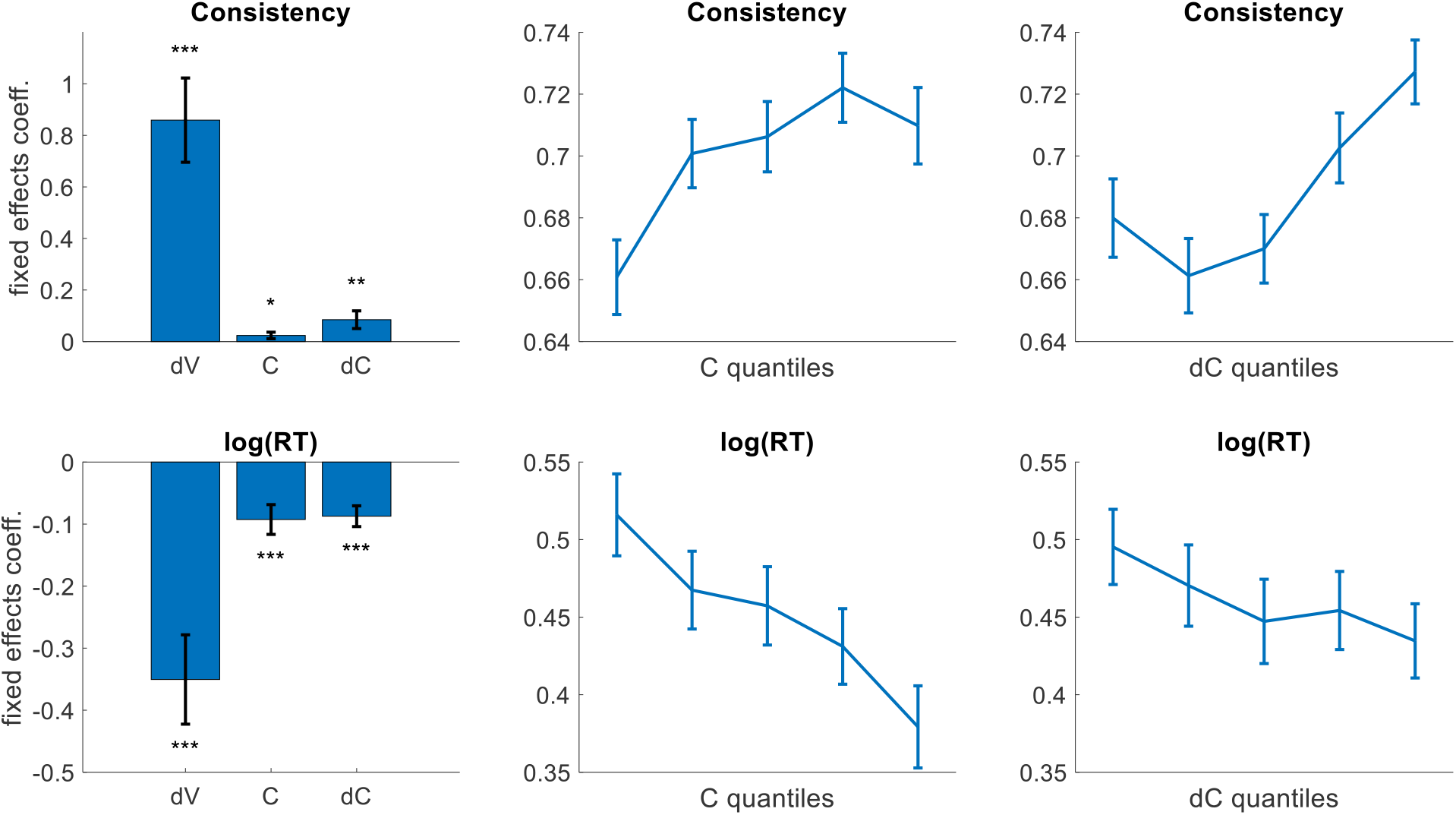
Previous experimental results (pooled across four studies, n=191) demonstrate the relationships between value difference (dV = V1 - V2), value certainty (C = C1 + C2), and certainty difference (dC = C1 - C2) with choice consistency and log(RT). We define V1 (V2) as the higher (lower) value rating, and C1 (C2) as the certainty report for the option with the higher (lower) value rating. The **upper** panels show the fixed effects coefficients from mixed effect logistic regression on choice consistency (left) and the isolated relationships between C and Consistency (center) and between dC and Consistency (right). The **lower** panels show the fixed effects coefficients from mixed effect linear regression on log(RT) (left) and the isolated relationships between C and log(RT) (center) and between dC and log(RT) (right). Error bars represent standard errors. Significance stars represent *** p < .001, ** p < .01, * p < .05 (one-sided t-tests). The data was binned by within-participant quantiles of C and of dC in the center and right plots, respectively. (data from Lee & Daunizeau, 2020, 2021; Lee & Coricelli, 2020; Gwinn & Krajbich, 2020)

A natural way in which one could introduce option-specific noise in the preferential DDM, consistent with the assumptions of the optimal policy model (Tajima et al, 2016; Fudenberg et al, 2018), would be to assume that the diffusion noise parameter increases with value uncertainty. This is because alternatives whose values are less certain will have greater variance in their representations, from which the evidence samples are drawn. However, on its own, this would lead to the wrong qualitative predictions with regard to the RT x certainty benchmark. Specifically, under such a model, diffusion noise would decrease as certainty increases, resulting in longer RT. Thus, unlike what we see in experimental data, such a model would predict that people would decide more quickly when they were less certain of the options’ values (all else equal). Moreover, this prediction is not specific to the classic DDM, but applies to the broader class of evidence accumulation-to-bound models (e.g., independent accumulators: Vickers, 1970; Brown & Heathcote, 2006; *leaky competing accumulator* or LCA: Usher & McClelland, 2001), which also predict that higher noise in the system will result in *faster* responses, in direct contrast to the empirical data^3^. An alternative way to include option-specific uncertainty in the DDM could be to assume that the response boundary increases with value uncertainty (while keeping the diffusion noise independent of value certainty). Such a model could account for the observed negative correlation between RT and certainty sum (C), but not certainty difference (dC; see Figure 2, bottom right panel). Furthermore, such a model would also predict that choice consistency (i.e., choices in favor of the higher rated options) decreases with certainty (resulting from a decrease in the response boundary, all else equal), which contradicts the empirical data. One possibility, which we will consider here (inspired by the DFT model) is that value uncertainty has a simultaneous effect *on both* the diffusion noise and the response boundary (each increasing with uncertainty; Busemeyer & Townsend, 1993; see Table 4, in particular). In this case, the impact of value certainty on choice accuracy would thus involve two opposing factors (noise and boundary), which may allow a way to account for some aspects of the data (although, note that this is not likely to provide an account for the dependency of choice consistency and RT on dC). Another hypothesis is that option-specific value certainty affects the DDM process through the drift rate. We provide preliminary support for this hypothesis in the Supplementary Material, using median splits on certainty and comparing the fitted model drift rate parameters.

The aim of this paper is to examine a number of DDM variants for preferential choice, to determine which can best account for the impact of value certainty on choice consistency and RT previously observed in experimental data (Lee & Daunizeau, 2020, 2021; Lee & Coricelli, 2020; Gwinn & Krajbich, 2020). In particular, we first present a variety of extensions of the classic DDM, each of which incorporates the concept of option-specific value certainty in a unique and realistic way. We start with the classic DDM (Model 1), which has no option-specific noise, and progress to models that include option-specific certainty in the sampling noise or in the response boundary (Models 2 and 3; although we know that these models are unlikely to succeed, we believe that it is instructive to formally test them and to examine their specific failures). We then examine a model (2+3) in which value certainty modulates both the sampling noise and response boundary, as well as models (4 and 5) based on signal-to-noise ratio concepts in the formulation of the drift rate. In the next section, we introduce formal notation for each of the models, and describe the models from a technical perspective. We then use simulations to demonstrate the theoretical predictions of each model, with respect to the impact of value certainty on choice consistency and RT. Finally, we fit each of the models to experimental data from four empirical datasets, quantitatively compare the performance of the models across the datasets, and use the results to suggest the best recommended approach for future studies to incorporate option-specific value certainty in models derived from the DDM. To anticipate our results, we find that a signal-to-noise DDM variant provides the best fit to the data and also meets all of the qualitative benchmarks.

## Computational Models

In each of the models described below, we consider decisions between two alternatives, with value estimates V1 and V2 (V1>V2) and value estimate certainties C1 and C2, respectively. Evidence of these values is integrated over deliberation time, subject to noise. The evidence accumulator for each decision is initialized at a neutral starting point (i.e., zero evidence or default bias in favor of either option), and evolves until reaching one of two symmetric boundaries (see Figure 1 above). For each decision, the output of the model is: the choice (ch = {0, 1}, where 1 indicates that the higher-valued option was chosen), determined by which boundary is reached; RT, determined by the number of integration time steps elapsed before that boundary is reached.

### Model 1: Classic DDM

As a baseline model for comparison (without any option-specific certainty term), we first consider the classic DDM. In this model, the equation that governs the evidence accumulation process is:

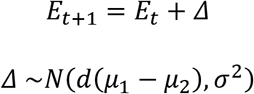

where *E* represents the cumulative balance of evidence that one option is better than the other, t represents the time from the start of deliberation, Δ represents the incremental evidence in favor of one option over the other at each time step*, d* is a sensitivity parameter, μ_i_ is the subjective value of option i, and σ^2^ represents processing noise in the evidence accumulation and/or comparator systems. In the classic DDM, choice probability and expected response time can be analytically calculated using the following equations (Alos-Ferrer, 2018):

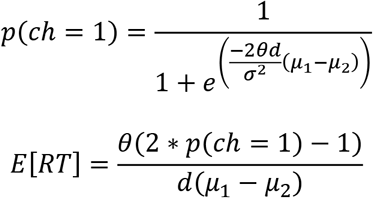

where θ is the height of the evidence accumulation boundary at which a decision is triggered (shown here as the upper boundary, for a choice of option 1), p(ch=1) is the probability that the upper boundary will be reached (rather than the lower boundary), and RT is the expected time at which the accumulation process will end (by reaching either of the boundaries) ^4^ . Choice probability and RT will thus be functions of the drift rate coefficient *d*, the diffusion (noise) coefficient σ^2^, and the height of the response boundary θ (see Figure 3)^5^. As expected, consistency increases and RT decreases with the drift rate. Increasing noise reduces both accuracy and RT, whereas increasing boundary increases both accuracy and RT.

**Figure 3:**
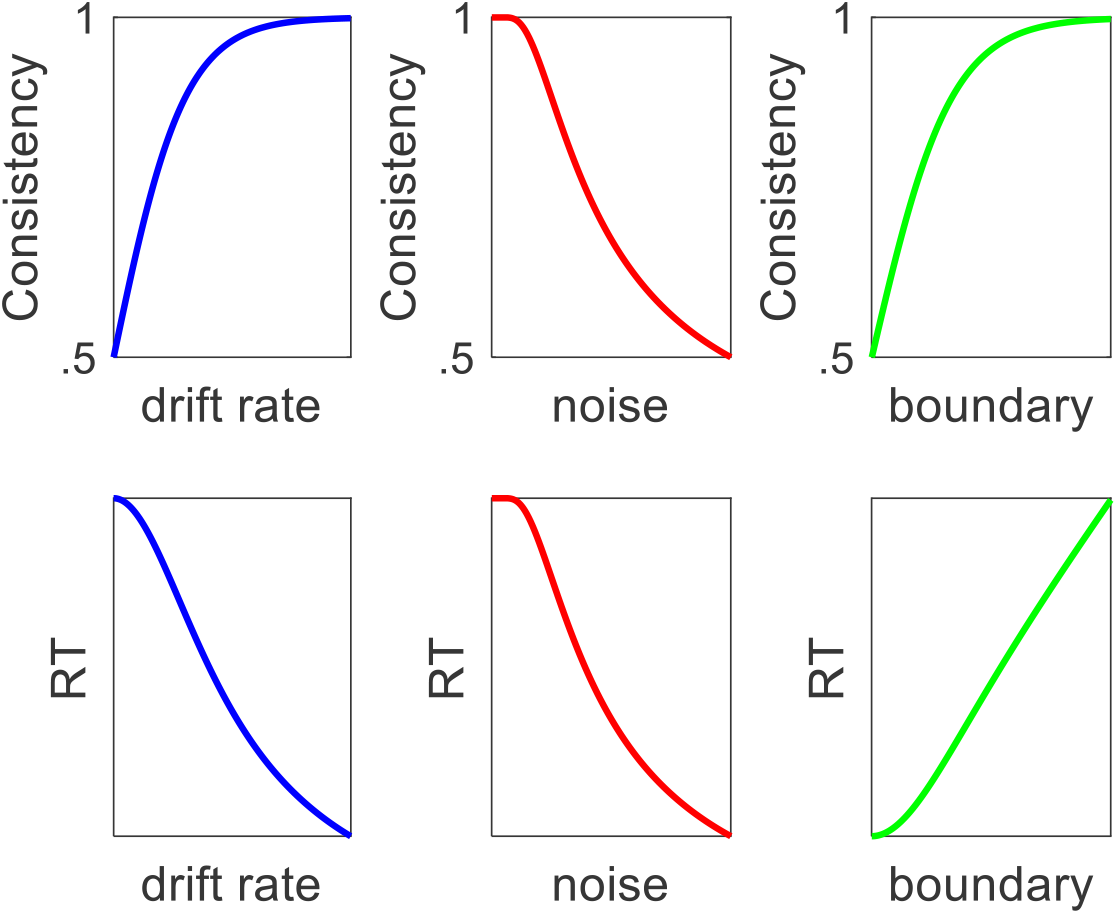
Dependency of choice consistency (upper row) and of RT (bottom row) on DDM parameters: drift rate (left/blue), diffusion noise (middle/red) and response boundary (right/green). With all other parameters fixed: an increase in drift rate leads to an increase in the probability of choosing the best option from 0.5 (random guess) to 1 and a decrease in RT; an increase in processing noise leads to a decrease in the probability of choosing the best option and a parallel decrease in RT; an increase in response boundary leads to an increase in the probability of choosing the best option and an increase in RT.

### Model 2: Certainty-Adjusted Diffusion Noise

One simple potential solution to incorporate option-specific uncertainty into the DDM would be to model the evidence accumulation process as:

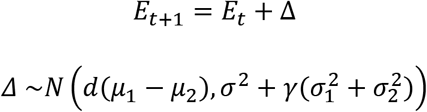

This corresponds to the classic DDM equation capturing the evolution of accumulated evidence across time, but with σ_i_^2^ representing the uncertainty about the value estimate of option i, and γ serving as a sensitivity parameter that controls the impact of this value (un)certainty on the level of diffusion noise. The only difference between this formulation and the classic one is that here the variance of Δ is specific to the options in the current choice set, whereas in the classic DDM, it is fixed across all choices (for an individual decision maker). A direct result of this reformulation is that choices between options with greater value uncertainty (lower C) will be more stochastic and take less time (on average), as can be seen by examining the (revised) DDM equations for choice probability and expected response time (see also Figure 3, center panels):

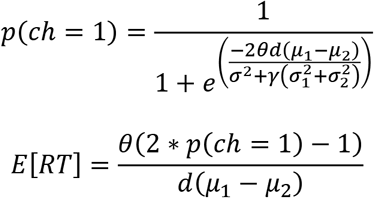

While the predicted relationship between certainty and choice consistency would match empirical observations, the relationship between certainty and RT would conflict with the empirical data.

### Model 3: Certainty-Adjusted Response Boundary

Another potential solution would be to allow the magnitude of the response boundary to vary as a function of option-specific (or trial-specific) value certainty. The evidence sampling process would be identical to that of the classic DDM. Under this model, the height of the boundary would increase as the value certainty of the pair of options decreased, on a trial-by-trial basis. Choice probability and mean RT would thus be calculated using the following equations:

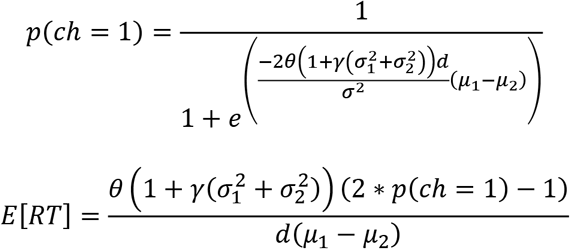

where γ is a sensitivity parameter that controls the impact of certainty on the magnitude of the response boundary. As can be seen in the equations, increasing value uncertainty (all else equal) will result in higher choice consistency and higher RT (see also Figure 3, right panels). While the predicted relationship between certainty and RT would match empirical observations, the relationship between certainty and choice consistency would conflict with the empirical data.

### Model 2+3: Certainty-Adjusted Diffusion Noise and Response Boundary

*Decision Field Theory* (DFT; see Table 4 in Townsend & Busemeyer, 1993) suggests that value certainty *simultaneously* modulates the diffusion noise and the response boundary. We therefore considered such a formulation in our set of models. The basic idea is that while uncertainty is expected to increase the sampling noise (see also Tajima et al, 2016), decision makers are likely to compensate for the reduced certainty by increasing their *caution parameter* – the response boundary. A version of DDM that adheres to this DFT principle would define the boundary as a function of the noise, which itself would be a function of value certainty. In this DFT-inspired model, choice probability and mean RT would thus be calculated using the following equations^6^:

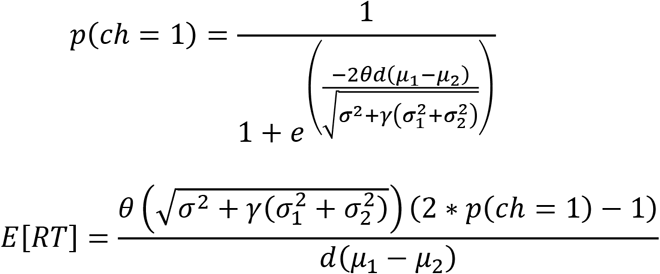

### Model 4: Certainty-Adjusted Drift Rate (value difference)

Another possible way in which the concept of option-specific value (un)certainty could be incorporated into the DDM would be through a signal-to-noise type dependency in the drift rate. The drift rate in the DDM symbolizes the momentary accumulation of evidence for one option over the other, equal to the value of one option minus the value of the other option (scaled by a fixed term). The accumulator variable is referred to as “evidence” because the probability distributions controlling it (or the neural activity driving it) are thought to provide a reliable signal that will accurately inform the decision. If the value representations of different options can have different levels of uncertainty, it stands to reason that the reliability of the “evidence” that these signals provide about the correct decision might also be different. As such, evidence drawn from a more reliable source (i.e., one with higher certainty) should be weighted more heavily. Under this framework, the equation governing the DDM process would be:

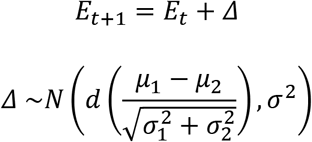

where σ^2^ (without a subscript) is the noise in the system unrelated to the specific choice options. The only difference between this formulation and the classic one is that here the mean of the option value difference is divided by its standard deviation 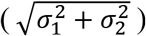. A direct result of this reformulation is that choices between options with greater value uncertainty will be less consistent and also take *more* time (on average), as can be seen by examining the (revised) DDM equations for choice probability and expected response time:

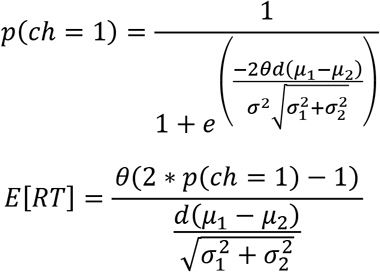

Here the impact of option-specific uncertainty on RT is more complex. First, greater uncertainty decreases RT through its effect on choice stochasticity (as before). Second, greater uncertainty directly increases RT by diminishing the slope of the drift rate. The second effect dominates. Note that Model 4 is mathematically similar to Model 2+3, because adjusting the drift rate in one direction is similar to simultaneously adjusting the response boundary and the diffusion noise in the other direction (as Model 2+3 does as a function of value certainty).^7^

### Model 5: Certainty-Adjusted Drift Rate (option values)

An alternative variant of the DDM in which the drift rate is altered by option-specific value certainty would be one in which the evidence in favor of each option *i* is scaled by its own precision term 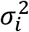, as is the case, for example, in multi-sensory integration (Drugowitsch et al, 2014; Fetsch et al, 2012). The drift rate would thus become the accumulation of adjusted evidence for one option over the other, equal to the precision-weighted value of one option minus the precision-weighted value of the other option (scaled by a fixed term). Here, the evidence drawn from a more reliable source (i.e., one with higher certainty) will be weighted more heavily (as in Model 4), but prior to a comparison between the alternative options. Note that here the certainty weighting is truly specific to each option, whereas in Model 4 the certainty weighting is specific to the pair of options. Under this framework, the equation governing the DDM process would be:

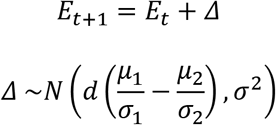

The only difference between this formulation and the classic one is that here the mean of the option value difference is adjusted by the standard deviations of the individual choice options. Because the evidence in favor of each option (prior to comparison) will be scaled by its own specific (and importantly, potentially different) precision term, the impact on both choice stochasticity and response time could go in either direction. This can be seen by examining the (revised) DDM equations for choice probability and expected response time:

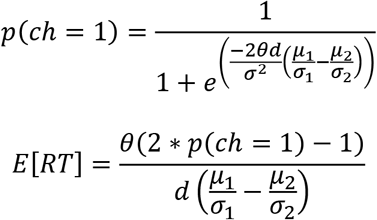

Here the impact of option-specific uncertainty on both choice and RT is more complex than in the previous models. If the value signal for one option has both a larger mean and a smaller variance, relative to the other option, the effective drift rate will be higher than in the classic DDM (e.g., choices will be less stochastic and faster). On the other hand, if the option with the larger mean value also has a higher variance (due to its value uncertainty), the effective drift rate might be lower than in the classic DDM (e.g., choices might be more stochastic and slower).

## Summary of Model Predictions with respect to the impact of value-certainty

We have considered a number of DDM variants, which make different predictions for how the value certainty of choice alternatives impacts the preference formation process, and in particular, how it affects choice consistency and RT. In Figure 4, we summarize these model predictions by simulating choice and RT output under each of the models, taking as input, value and certainty ratings generated from uniform distributions covering the full range available in the empirical data, without any correlation between the variables^8^, and using normally-distributed parameters with moments to match those in the best fits to the empirical data (see below for a description of the empirical data).

**Figure 4:**
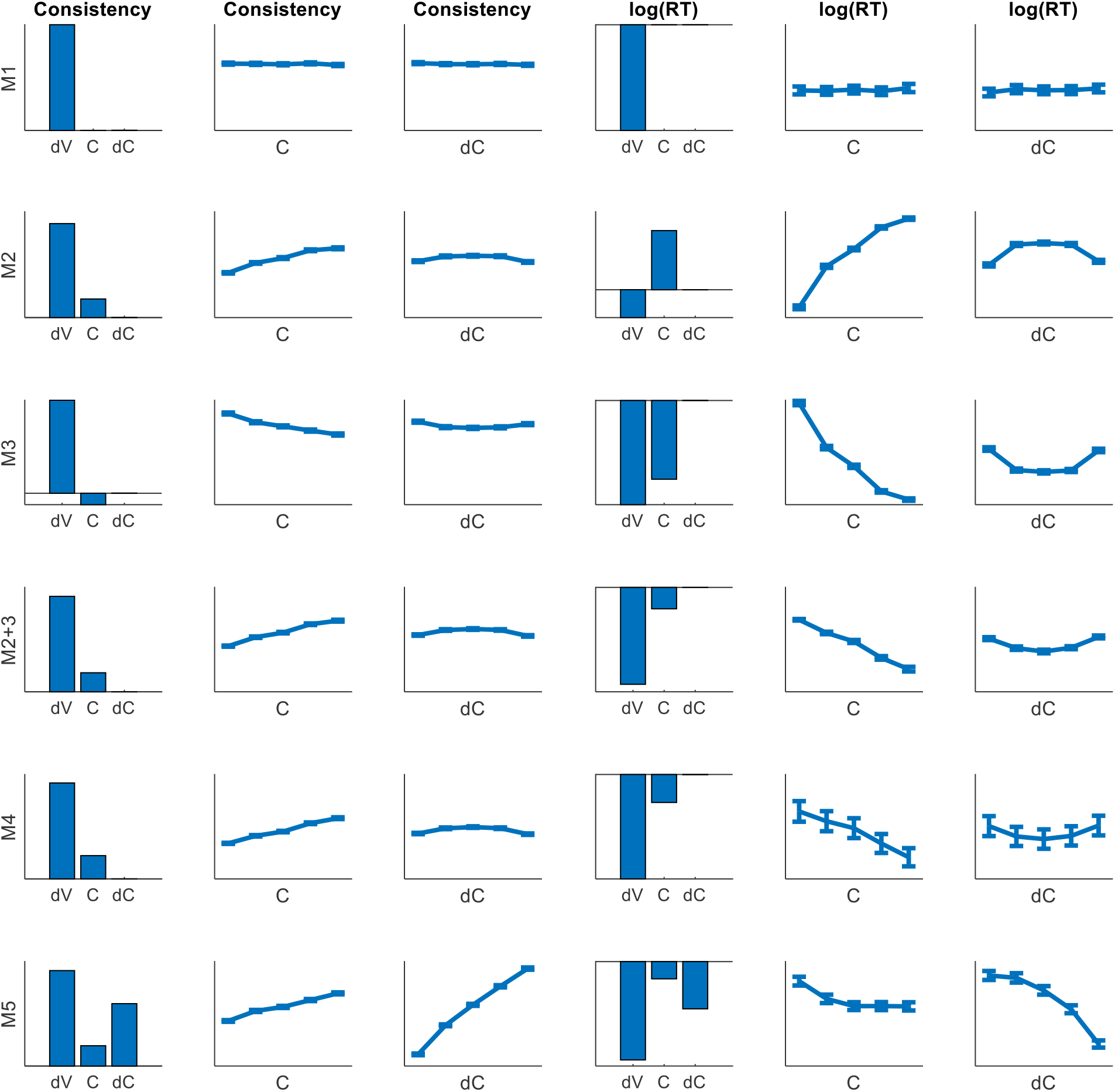
Theoretical model predictions for the impact of C and dC on consistency and on RT (data simulated using uniformly distributed (0:1] value estimates and certainty and normally distributed parameters with moments to match the experimental data).

As we can see, Model 1 (the classical DDM), which has no C-dependency in its evidence accumulation, shows no C-dependency in either choice consistency or RT. Model 2 makes the counterintuitive prediction that RT increases with value certainty. Model 3 makes the counterintuitive prediction that choice consistency decrease with certainty. Each of the other models predicts patterns more similar to the empirical data with respect to C, although only Model 5 appears to capture the effects related to dC.

### Empirical data

To test the performance of the set of models we introduced above against actual preferential choice data, we gathered datasets from four experiments reported in the literature that examined choices between pairs of food items, whose ratings (value and certainty) were measured for each participant before the pairs were presented for binary choice.

#### Dataset 1

The first dataset we examined was from Lee & Daunizeau (2020). In this study, participants made choices between various snack food options based on their personal subjective preferences. Value estimates for each option were provided in a separate rating task prior to the choice task. Participants used a slider scale to respond to the question, “Does this please you?” After each rating, participants used a separate slider scale to respond to the question, “Are you sure?” This provided a measure of value estimate certainty for each item. During the choice task, participants were presented with pairs of snack food images and asked, “What do you prefer?” After each choice, participants used a slider scale to respond to the question, “Are you sure about your choice?” to provide a subjective report of choice confidence. This dataset contained 51 subjects, each of whom were faced with 54 choice trials.

#### Dataset 2

The second dataset we examined was from Lee & Daunizeau (2021). In this study, participants made choices between various snack food options based on their personal subjective preferences. Value estimates for each option were provided in a separate rating task prior to the choice task. Participants used a slider scale to respond to the question, “How much do you like this item?” After each rating, participants used the same slider scale to respond to the question, “How certain are you about the item’s value?” by indicating a zone in which they believed the value of the item surely fell. This provided a measure of value estimate certainty for each item. During the choice task, participants were presented with pairs of snack food images and asked, “Which do you prefer?” After each choice, participants used a slider scale to respond to the question, “Are you sure about your choice?” to provide a subjective report of choice confidence. This dataset contained 32 subjects, each of whom were faced with 74 choice trials.

#### Dataset 3

The third dataset we examined was from Lee & Coricelli (2020). In this study, participants made choices between various snack food options based on their personal subjective preferences. Value estimates for each option were provided in a separate rating task prior to the choice task. Participants used a slider scale to respond to the question, “How pleased would you be to eat this?” After each rating, participants used a six-point descriptive scale to respond to the question, “How sure are you about that?” This provided a measure of value estimate certainty for each item. During the choice task, participants were presented with pairs of snack food images and asked, “Which would you prefer to eat?” After each choice, participants used a slider scale to respond to the question, “How sure are you about your choice?” to provide a subjective report of choice confidence. This dataset contained 47 subjects, each of whom were faced with 55 choice trials.

#### Dataset 4

The fourth dataset we examined was from Gwinn & Krajbich (2020). In this study, participants made choices between various snack food options based on their personal subjective preferences. Value estimates for each option were provided in a separate rating task prior to the choice task. Participants used a 10-point numerical scale to respond to the prompt, “Please indicate how much you want to eat this item.” After each rating, participants used a seven-point numerical scale to respond to the prompt, “Please indicate how confident you are in your rating of this item.” This provided a measure of value estimate certainty for each item. During the choice task, participants were presented with pairs of snack food images and instructed to choose the one that they preferred to eat. Choice confidence was not measured in this study. This dataset contained 36 subjects, each of whom were faced with 200 choice trials.

#### Computational Model-Fitting Procedure

We fit the experimental data to each of the models that we described above. We then performed Bayesian model comparison to determine which of the models (if any) performed better than the others across the population of participants. For this model fitting and comparison exercise, we relied on the Variational Bayesian Analysis toolbox (VBA, available freely at https://mbb-team.github.io/VBA-toolbox/; Daunizeau, Adam, & Rigoux, 2014) with Matlab R2020a. Within participant and across trials, we entered the experimental variables {left option value, right option value, left option certainty, right option certainty} as input and {choice = 1 for left option, 0 for right option; RT} as output. We took the inverse of the value certainty rating as our measure of variance for the value of each option. We also provided the model-specific mappings from input to output as outlined in the analytical formulas above. The parameters to be fitted included all of the *d*, σ^2^, γ, λ, and α terms described above in the model formulations. VBA requires prior estimates for the free parameters, for which we set the mean equal to one and the variance equal to *e* (to allow an arbitrarily-large step size during the gradient descent search algorithm, yet constrain the algorithm to a reasonable search space) for each parameter. VBA then recovers an approximation to both the posterior density on unknown variables and the model evidence (which is used for model comparison). We used the VBA_NLStateSpaceModel function to fit the data for each participant individually, followed by the VBA_groupBMC function to compare the results of the model fitting across models for the full group of participants.

The benefit of using VBA to fit the data to our models spawns from the nature of the preferential choice data itself, where the value estimate and certainty of each option necessarily varies from one trial to the next (i.e., no two choice trials are ever alike). Other DDM parameter estimation approaches that are popular in the literature (e.g, Voss & Voss, 2008) do not optimally handle this kind of experimental design, because they lack the trial repetitions per condition (in this case, value and certainty ratings) necessary to provide empirical estimates of RT distributions (or moments) in each condition. The VBA toolbox is computationally efficient, as it relies on Variational Bayesian analysis under the Laplace approximation. This iterative algorithm provides a free-energy approximation for the model evidence, which represents a natural trade-off between model accuracy (goodness of fit, or log likelihood) and complexity (degrees of freedom, or KL divergence between priors and fitted parameter estimates; see Friston et al., 2007; Penny, 2012). Additionally, the algorithm provides an estimate of the posterior density over the model free parameters, starting with Gaussian priors. Individual log model evidence scores are then provided as input to the group-level random-effect Bayesian model selection (BMS) procedure. BMS provide an exceedance probability that measures how likely it is that a given model is more frequently implemented, relative to all other models under consideration, in the population from which participants were drawn (Rigoux et al., 2014; Stephan et al., 2009). This approach to fitting and comparing variants of DDM has already been successfully demonstrated in previous studies (Lopez-Persem, Domenech, & Pessiglione, 2016; Feltgen & Daunizeau, 2021).

As an initial check to verify that our model-fitting procedure is suitable for this specific analysis, we performed a test of model recovery. Specifically, we created synthetic data for each participant in Studies 1-4, under two separate scenarios: 1) taking as input uniformly distributed data (value estimate and certainty for each option, uncorrelated) covering the same scale as the experimental data; 2) taking as input the actual rating data provided by each participant. We simulated choices and RT for each participant, separately according to each of the models and using the specific best-fit parameters for each participant for each model. We then fit the simulated data (per participant) to each of the models and performed the same formal model comparison as with our real experimental data. The results of this procedure can be seen below as model confusion matrices (see Figure 5). The matrices show, for each true generative model, the percentage of simulated participants (under that model) that were attributed to each of the best fit models by our model-fitting procedure. As shown in the first confusion matrix (Figure 5a), overall confusion was low when the input data was theoretically ideal, as the procedure attributed the true model as the best fit model for the vast majority of participants. Model 4 was the only model in our set of models that our model-fitting procedure was unable to accurately recognize (using ideal input data), as it wrongly classified about half of the participants simulated under Model 4 as having been simulated under Model 2+3. This is not surprising, since Models 2+3 and 4 are mathematically similar (see above). This exercise verified that our model-fitting procedure was theoretically capable of accurately distinguishing data produced by the different models, which suggests that our quantitative model comparison based on actual empirical data (see below) should be valid. As shown in the second confusion matrix (Figure 5b), the models were also recovered (though less accurately) when using the actual experimental input data. Some confusions arose for about half of the participants simulated under Model 2, who were wrongly attributed to Model 1 (due to its reliance on fewer free parameters). Also Model 2+3 stole a substantial amount of support from Model 4. Nevertheless, even when the input data was less than ideal, overall average model recovery (52.5%) was well beyond chance level (16.7%).

**Figure 5:**
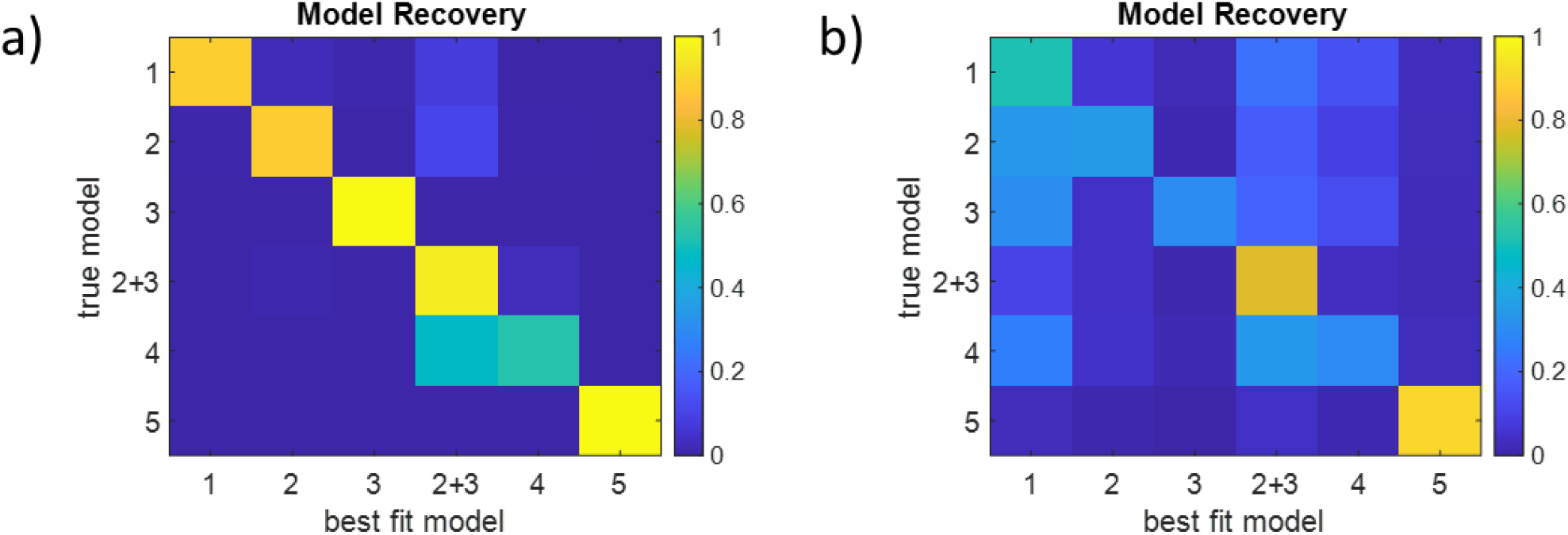
Model recovery analysis. Each cell in the “confusion matrix” summarizes the percentage of simulated participants (under each true model) for which our model-fitting procedure attributed each of the (best fit) models. a) Uniformly distributed value and certainty input data. b) actual experimental value and certainty input data.

#### Quantitative Model Comparison

The classic DDM, our Model 1, has been validated countless times for its ability to account for two-alternative forced choice responses and mean response times. The other models described above are new and have therefore never been tested with empirical data. We thus performed a formal model comparison of our full set of models, with Model 1 serving as a benchmark. The results are displayed in Figure 6, which shows the support that each model received within participants (left panel) and across participants (right panel). We provide a summary of the best-fitting parameters for each model in the Supplementary Material.

**Figure 6:**
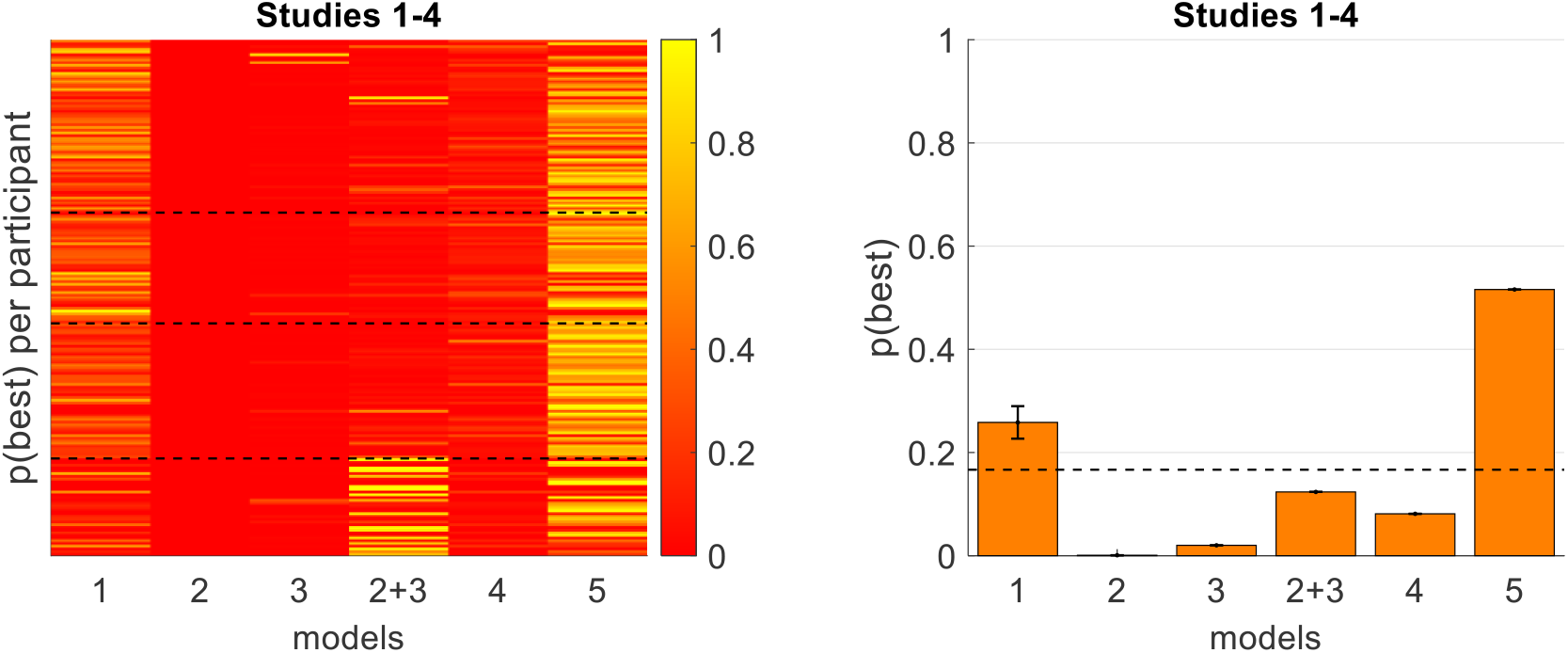
Model comparison results across all core models. We show here the probability that each model best accounted for the data at the participant level (left panel), across the four studies we examined; each cell represents the probability that the model (column) best represents the behavior of the participant (row). The dashed lines serve to indicate which participants belonged to each of the four datasets. We also show the probability that each model best explains the data across the participant population (right panel), across all studies. The dashed line indicates chance level if all models were equally probable a priori.

Across all four studies, Models 2 and 3 were each dominated by Model 1 (the classic DDM) as anticipated, as were Models 2+3 and Model 4. Model 5, by contrast, dominated Model 1 (estimated cross-participant model frequency of 0.516 for Model 5 versus 0.258 for Model 1). In recognition of the possibility that the impact of certainty on the drift rate might be non-linear, we considered a variation of our best-fitting model (Model 5), which we label Model 5*. Here, the only change was the inclusion of an exponent α in computation of the signal to noise ratio:

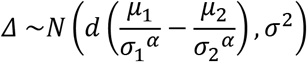

In a formal comparison of Model 5 against Model 5*, Model 5 was the clear winner with an estimated model frequency (across participants) of 0.670 (see Figure 7). It appears that although the flexibility associated with Model 5* improves the fit for some individual participants, at the group level, the added flexibility offered by the exponent parameter was not great enough to offset the complexity cost for adding an extra parameter.^9^

**Figure 7:**
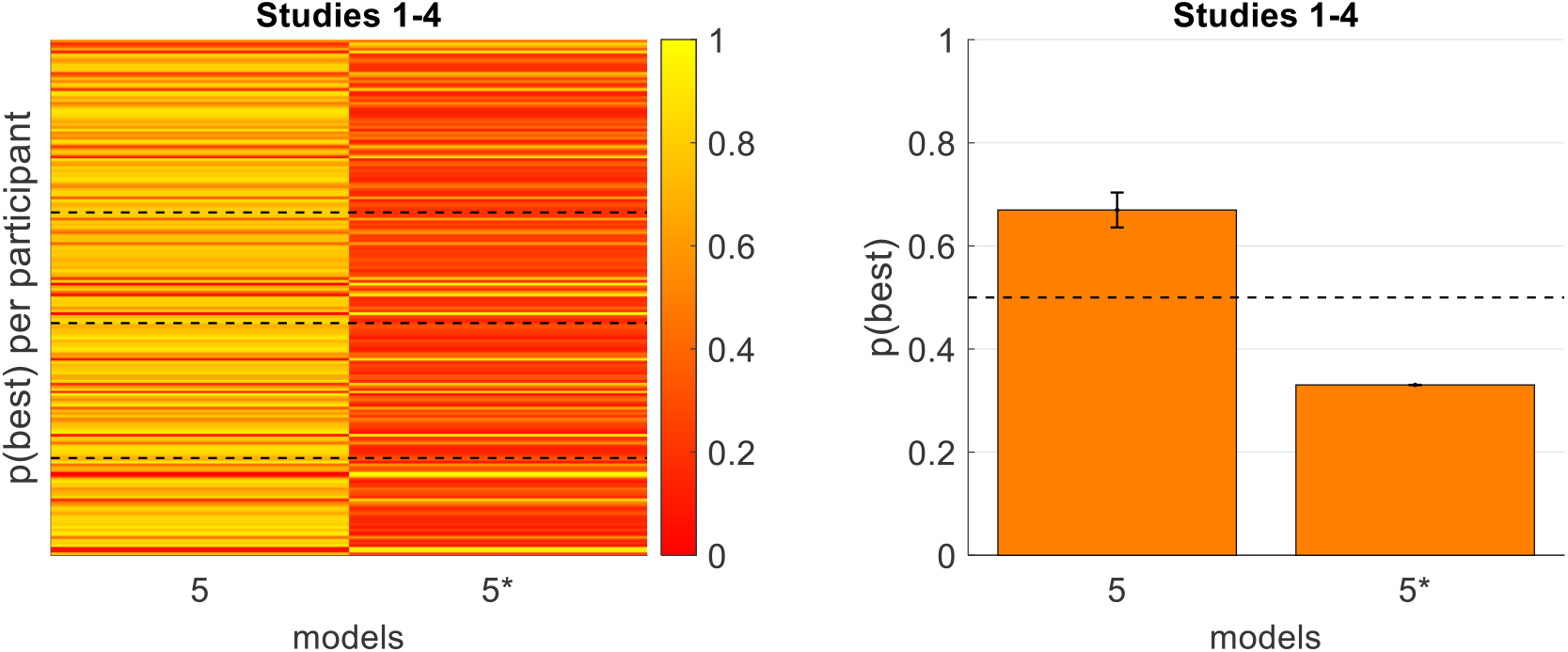
Model comparison results for Model 5 vs Model 5* (same format as Figure 6).

To summarize, the quantitative model comparison of the DDM variants favors the signal-to-noise variant (Model 5). To better understand why Model 5 wins the quantitative model comparison, it is useful to examine how the various models account for the qualitative relationships between certainty and choice consistency or RT in the data (Figure 2). Towards this aim, we used the actual ratings provided by each participant and the best fitting model parameters (for each participant under each model) to generate synthetic data for each combination of participant and model. We then carried out the same procedure that we applied to the empirical data (Figure 2) to the data simulated under each of the models. These results are shown in Figure 8 (the first row shows the empirical data and the following rows show the data simulated under each of the models).

**Figure 8:**
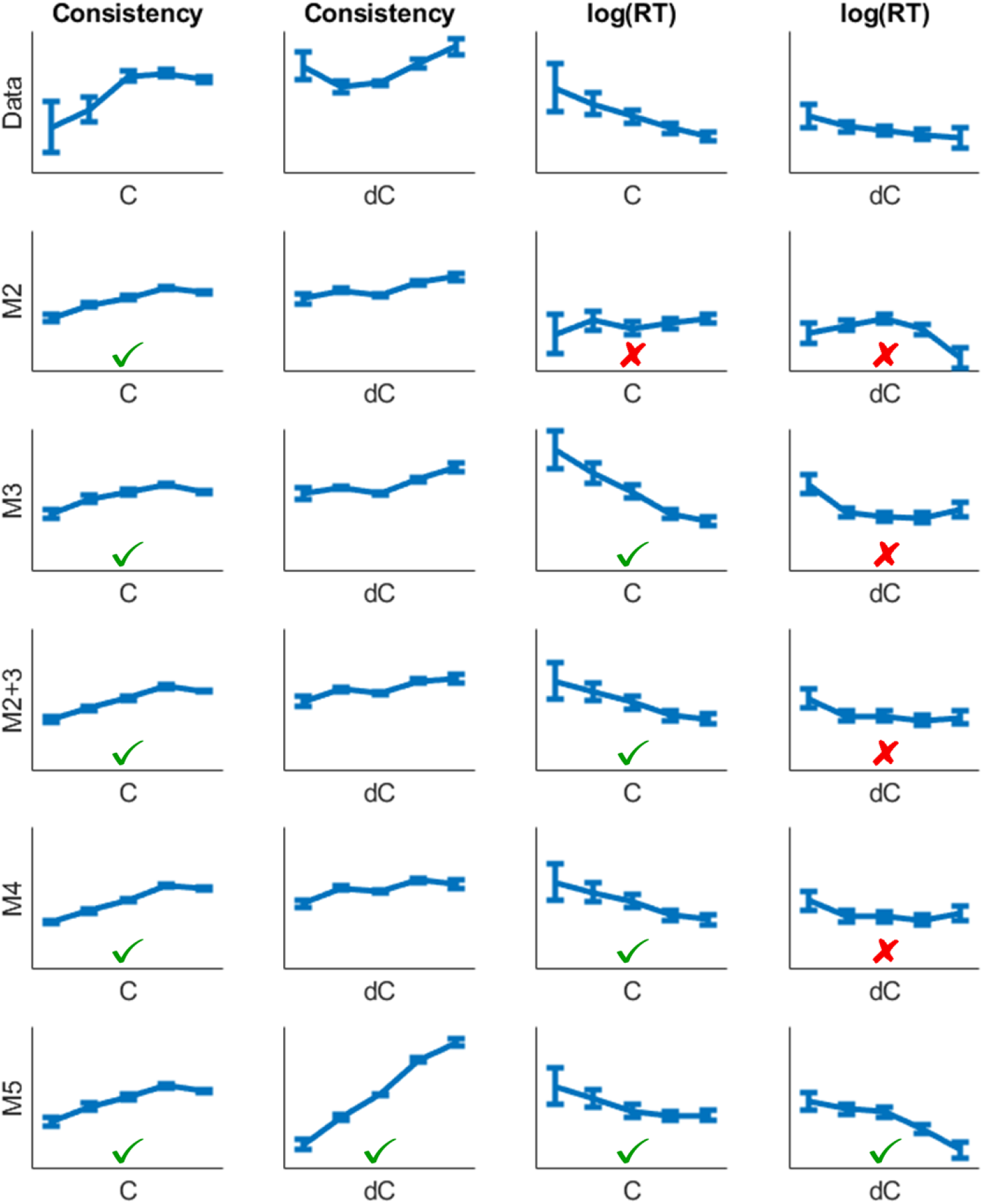
Qualitative predictions of the effects of value difference (dV = V1 - V2), value certainty (C = C1 + C2), and certainty difference (dC = C1 - C2) on choice consistency and log(RT) for simulated data for each model; the top row shows the empirical data, while each subsequent row shows data simulated under each of the models using the actual input and best-fitting parameters for each participant; the curves represent within-bin means across participants; the data is separated by uniform bins covering the full possible range of C (0,2] and dC (−1,1); error bars represent s.e.m. Green checkmarks indicate a qualitative match between model predictions and empirical data; red Xs indicate a lack of qualitative match.

In order to simplify the comparison between the data and the models, we have included binary indicators that correspond to the match between the data and the model dependency shown (green checkmark for a good match, red X for a poor match)^10^. As we can see, Model 5 seems to be the model best able to account for the qualitative relationships between certainty and the choice variables (see in particular the dependency of RT on dC). As we show in the Supplement this model also accounts well for the dependency of the shape of the RT distribution on model predicted drift-rate (Fig S2).

## DISCUSSION

The aim of this study was to examine a number of extensions of the drift-diffusion model for preferential choice that include option-specific value certainty, and to probe them in their ability to account for benchmark data on the dependency of choice consistency and RT on value certainty. As illustrated in Figure 2, the experimental data that we examined show that certainty has a clear and systematic impact on both consistency and RT, thereby motivating an extension of the classic DDM (with no option-specific uncertainty component) and providing constraints on the way one can introduce option-specific uncertainty into the models. As we have shown, perhaps the simplest DDM extension, in which (only) the evidence accumulation noise decreases with value certainty, produces the wrong qualitative prediction: RT increases with certainty (as certainty reduces the stochasticity of the system, which slows down RT; see Figure 3, center panels). Moreover, the problem with this method of introducing option-specific value certainty in modelling value-based decisions is not particular to the DDM, but also applies to the broader class of evidence accumulation-to-bound models (e.g., LCA, Usher & McClelland, 2001; independent accumulators, Vickers, 1970), in which noise decreases RT.

An alternative way to introduce option-specific value uncertainty in the DDM framework could be to assume that the uncertainty affects (only) the response boundary. Accordingly, decision makers would compensate for their uncertainty by increasing the height of the boundary. While such a model could account for the negative correlation between RT and certainty (C; see Figure 2), it would not be able to account for the negative correlation between RT and certainty difference (dC). Moreover, such a model would predict that choices become more stochastic as value certainty increases (due to more random crossings of the lower boundary; see Figure 3, right panels), which is both counterintuitive and in contrast to the experimental data (see Figure 4). We also considered a model in which both diffusion noise and response boundary were simultaneously modulated by value certainty, which can make the correct predictions in terms of C. However, this model does not predict any impact of dC on either choice consistency or RT (see Figure 4). Thus, we believe that the way in which value uncertainty affects the decision process is via its impact on the drift rate. (We provided additional support for this conclusion in the Supplementary Material – Figure S8 – using median splits on certainty and comparing the fitted model parameters.)

In order to understand the way in which value certainty affects the drift rate, we examined and tested two core DDM variants built on certainty-modulated drift rates. These models (4 and 5) were based on signal-to-noise principles. While Model 4 was able to account for some of the qualitative relationships in the data, only Model 5 accounted for all of them.^11^ In this model, the drift rate of the diffusion process is not simply the fluctuating difference in the values of the options (Tajima et al, 2016), but rather a difference between the ratios of the mean values and their corresponding value uncertainties. This mechanism has a normative flavor, as it penalizes values that are associated with uncertain alternatives. Some similar types of signal-to-noise models have also been supported by data in perceptual choice tasks. For example, de Gardelle and Summerfield (2011) examined choices in which an array of eight visual patches of variable color or shape are compared (in binary choice) to a reference (color or shape). By independently varying the set mean distance (in the relevant dimension) from the reference as well as the set variance, they found that both independently affect choice accuracy and RT. In particular, the set variance (which is the analog of our value uncertainty) reduces choice accuracy and increases RT. As shown by de Gardelle and Summerfield (2011), a signal-to-noise model can account for this dependency. Furthermore, the random dot motion task that is widely used alongside the DDM in perceptual decision-making studies provides a signal-to-noise ratio as input for the drift rate (e.g., Gold & Shadlen, 2007); for this task, drift rate is typically determined by the motion coherence, which is composed of the number of dots moving in the same direction (signal) as well as the number of dots moving randomly (noise).

We showed via simulation that only Model 5 can account for all of the qualitative relationships observed in the empirical data between certainty sum and certainty difference, and choice consistency and mean RT (see Figures 4 and 8). Under Model 2, increased certainty leads to decreased diffusion noise. Although this would appropriately lead to more accurate responses, it would paradoxically lead to slower RT. Under Model 3, increased certainty leads to a lower response boundary. Although this would appropriately lead to faster responses, it would paradoxically lead to less consistent responses. Under Model 2+3, inspired by DFT principles, increased certainty leads to both decreased diffusion noise *and* a lower response boundary. This model thus balances the two parameters against each other and is able to predict that higher certainty leads to both faster and more accurate responses. However, it is agnostic to the effects of certainty difference, which were prominent in the data. Under Model 4, increased certainty leads to an increased drift rate (note that this is mathematically similar to a decrease in both diffusion noise and response boundary, as the DDM is over-specified by these three parameters). This would appropriately lead to both faster and more accurate responses. However, Model 4 cannot account for effects related to certainty difference, because certainty only enters this model in relation to the choice pair and not the individual options. Only Model 5 successfully predicts all effects related to both certainty and certainty difference. This model is similar to Model 4, in that certainty positively impacts choice consistency and negatively impacts RT through its modulation of the drift rate. However, unlike the other models, certainty enters Model 5 in relation to each option independently. It is for this reason that Model 5 is also able to predict the effects related to certainty difference (see the Supplementary Material for more details).

Future work will be needed to examine the neural mechanism that extracts the drift rate from fluctuating values (sampled from memory or prospective imagination; Bakkour et al, 2019; Poldrack et al, 2001; Schacter, Addis, & Buckner, 2007) and that reduces the drift rate of more strongly fluctuating items. One interesting possibility is that the effective drift rate might be modulated by the temporal congruency of the evidence in successive samples (Glickman, Moran, & Usher, 2020); the higher the value certainty, the lower the variance of the value signals, thus leading to a higher probability that successive samples will provide consistent choice evidence. Perhaps certainty could be tuned by attentional gating in the brain valuation pathways (Schonberg & Katz, 2020), such that more attention to the valuation process would enable higher value certainty. Future research is also needed to examine if the effects of value uncertainty on choice correlate with risk- or ambiguity-aversion at the level of individual participants, and to integrate this type of model with dynamical attentional affects as in the attentional drift-diffusion model (aDDM; Krajbich et al, 2010; Sepulveda et al, 2020). Recent work has suggested that attention allocated toward a particular choice option will serve to reduce the uncertainty about the value of that option (Callaway, Rangel, & Griffiths, 2020; Jang, Sharma, & Drugowitsch, 2020). If true, this would provide a link between the current work and the “gaze bias” effect (Krajbich, Armel, & Rangel, 2010; Sepulveda et al, 2020).

### Response Time Distributions

One of the strengths of DDM-type models is their ability to account not only for accuracy and mean RT, but also for how the full distribution of RT varies with task conditions (see Ratcliff & McKoon, 2008, for a review). In this study, we did not fit RT distributions for a pragmatic reason: the preference data (Studies 1-4) do not have enough repeated trials per condition (i.e., specific combinations of value and certainty rating) needed to estimate RT distributions. Nevertheless, we thought to examine predictions for how RT distributions vary with the predicted drift rate (which is a function of the value estimates, V1 and V2, and their certainties, C1 and C2) in the signal-to-noise model. As we report in the Supplementary Material (Figure S2), we find the typical rightward-skewed RT distributions in data simulated under the snDDM. Moreover, we find the expected patterns with respect to trials with low versus high predicted drift rate. The experimental data appear to confirm the model predictions (see Figure S2). This suggests that the snDDM can account for patterns in the full RT distributions, in addition to all of the qualitative and quantitative support that we reported above. However, future studies that include more trials per condition (i.e., each specific level of dV, C, and dC) will be needed to fully validate this.

### Impact of Overall Value on Response Time

In this study, we focused on the impact of value difference, certainty sum, and certainty difference on choice behavior. Yet there remains an additional element that we left out: value sum (V = V1 + V2). A number of previous studies have exposed a clear and robust effect of overall value (of the options being considered during a particular decision) on RT, where responses are typically faster for higher-valued options (Smith & Krajbich, 2019; Polania et al, 2014; Hunt et al, 2012; Teodorescu, Moran & Usher, 2016; Sepulevada et al, 2020). We also found this effect in the datasets we examined (see the Supplementary Material). As shown in Figure S5, the value sum (V) has a clear effect on RT (though somewhat smaller than the effect of dV). People decide faster (all else equal) on a choice between a pair of alternatives with ratings 0.9 and 0.8, respectively, than on a choice between alternatives with ratings of 0.2 and 0.1, respectively. This overall value effect is naturally explained in race or competing accumulator models (Vickers, 1970). It is also explained in the LCA (Teodorescu, Moran & Usher, 2016) or DFT^12^ models. However, the overall value effect is more challenging to explain within a diffusion model, in which the preference formation is based on relative evidence and unrelated to summed overall evidence. Interestingly, although not predicted and therefore not discussed so far, it turns out that the snDDM can account for the negative relationship between value sum and RT (see Figure S5). The snDDM account of this effect is an emergent one (i.e., there was nothing in the model that was explicitly dependent on V). We briefly comment on this in the Supplementary Material, and leave it to future studies to explore this effect further.

### Relation to Other Models and Additional DDM Variants

The most influential model for preferential choice is the *decision field theory* (DFT; Busemeyer & Townsend, 1993; Busemeyer & Diederich, 2002; Busemeyer et al, 2019; Roe et al., 2001). The DFT model was the first model that adopted the sequential sampling (integration to bound) concept in its account of preferential choice. The model was applied to risky choice (Busemeyer & Townsend, 1993), where it was able to account for violations of the independence axiom (the Myers effect; Busemeyer & Townsend, 1993, Fig. 2), and to multi-attribute decisions (Roe et al., 2001), where it accounted for contextual reversal effects (the attraction, similarity, and compromise effects; see also Johnson & Busemeyer, 2005, for an account of preference reversals between choice and price elicitation in risky choice). Moreover, the DFT proposed a mechanism for the stochastic nature of the evidence accumulation process, which it related to fluctuations in attention between event outcomes or decision attributes. The DFT model predicts that the magnitude of this stochastic component (the diffusion noise term) increases with the outcome variance associated with the options (which in turn reflects uncertainty). While in this study we do not focus on risky choice nor on explicit multi-attribute choices (we focus here on holistic and potentially memory-based choices between food items), and we only consider relatively simple DDM variants, it is important to discuss how our results compare with DFT-inspired uncertainty modulations.

As we described above, the DFT suggests that uncertainty has a dual effect on the preference formation — it increases both the diffusion noise and the response boundary (as a type of compensation for the former). Our Model 2+3 version of the DDM was based on this idea, and it successfully accounted for the qualitative effect of certainty sum (C) on choice consistency and on RT. This model could not, however, account for the effect of certainty difference (dC; the certainty of the high-value option minus the certainty of the low-value option) on consistency or on RT. This DDM extension was therefore outperformed by the snDDM in its quantitative account of the data. Obviously, this is not a test of the DFT model (which is more complex, including other factors such as an information leak, and which has not yet been applied to the type of tasks we examined here), but merely suggests that value certainty is more likely to operate (in our task) via a direct effect on the drift rate (rather than via modulation of the diffusion noise and of the response boundary). Future work will be needed to examine in detail how the DFT accounts for choice and RT for this type of preferential choice data.

We have focused here on simple DDM variants that include fixed (i.e., static across decision time) response boundaries. While this is the type of DDM that has been explored most in experimental psychology (Ratcliff, 1978; Ratcliff & Rouder, 1998), other contemporary versions of the DDM sometimes include response boundaries that collapse over time (Evans, Trueblood, & Holmes, 2020; Hawkins et al, 2015; Palestro et al, 2018; Glickman & Usher, 2019; Glickman et al, 2019). Such collapsing bounds were first introduced to truncate the long tails of the RT distributions that are predicted by a fixed-boundary DDM, especially for error trials (Milosavljevic et al, 2010). Some researchers have since included some form of collapsing bound parameter in their usage of the DDM, because it tends to provide better fits to the data (for preferential choice see Glickman et al, 2019). Furthermore, Tajima et al (2016) and Fudenberg (2018) proved that the DDM was an optimal model only in the case that the response boundary decreased across deliberation time (although their work related to series of sequential decisions that created an explicit opportunity cost of time, thereby mandating the collapsing bounds). In the current study, we remained agnostic as to the shape of the DDM response boundary (fixed versus collapsing). Whereas we relied on the fixed-boundary DDM in our model comparison, we verified that the inclusion of a collapsing boundary does not change any of our conclusions (see Figure S1A in the Supplementary Material), showing that, like for the fixed-boundary DDM (see Figure 3, center panel), RT in the collapsing-boundary model also decreases with sampling noise that could be associated with value uncertainty (contrary to the data; see Figure 2). This can easily be understood by examining the basic relationships between sampling noise and both response probability and RT. If there were no noise in the sampling/accumulation process, the evidence would simply accumulate along a straight trajectory until reaching a bound. Adding noise perturbs the trajectory (at any point in time) and thus increases the probability that the response boundary will be reached sooner. This is because the probability of reaching a boundary on the next time step is a decreasing function of the current distance from the accumulator to the boundary, and an increasing function of both the mean and the *variance* of the trajectory (which reflects the accumulation noise). Note that this will hold for any response boundary, regardless of the shape. Thus, models such as our Model 2 will also fail even when including collapsing boundaries (see Figure S1A in the Supplementary Material for a confirmation of this via simulations).

Some versions of the DDM (Ratcliff & Rouder, 1998) include additional parameters, which we did not examine in this study. Such parameters include non-decision time (time for encoding and response, or T_er_), starting point bias (and starting point variability), and drift rate variability. While it might be mathematically possible to account for the effects of option-specific certainty by allowing T_er_ to vary as a function of certainty, we believe that this would not be a principled approach. Because T_er_ corresponds to time spent on perceptual (encoding) or motor (response) processes, we see no reason why this should vary with value certainty (determined by valuation processes). Similarly, we did not consider a starting point bias in the current study, as such a bias is typically meant to capture prior expectations about which option is more likely to be better. In the experimental paradigms that we considered, there are no such prior expectations – the choice pairs were randomly created, each option was randomly assigned as either option 1 or option 2, and the sequence of trials was randomized. Therefore, as participants proceeded to make choices across trials, any potential biasing effects from one trial should be obliterated by the effects from all other trials, leaving the prior expectations about which option will be better (on the next trial) centered on zero. It remains possible that participants might have an inherent preference to pick the option displayed on the left (or right) side of the screen, regardless of content. However, we checked the data and found no evidence for a response side bias (see the Supplementary Material for details). It would also be theoretically reasonable to consider starting point variability (around a mean of zero). This could be due to a carry-over effect of the previous trial, where residual neural activation might lead the decision maker to be inclined to repeat the same choice on the next trial. However, this implies that starting point variability is caused by information from previous trials, and not from the current trial (for which deliberation will have not yet begun). Hence, the starting point variability is unlikely to be related to the option-specific value certainty on the current trial and therefore cannot mediate the impact of certainty on either consistency or RT. However, see the Supplementary Material for a confirmation (via simulations) that even if one were to imagine that starting point variability increased as a function of value uncertainty (a type of noisier responding), it would have no impact on the mean RT (Figure S1C), and it could definitely not account for the slowdown with value uncertainty shown in the data (Figure 2). Finally, we have not considered drift rate variability in the current study. In some versions of the DDM (see Ratcliff & McKoon, 2008), the drift rate on a given trial is a random variable selected from a normal distribution whose mean equals the value difference of the options on that trial and whose variance corresponds to either attentional or memory biases specific to that trial (meaning that if the same decision is presented again, the drift may differ). In principle, it is possible (but less likely, we think) that value uncertainty could affect the drift rate variability (across trials) rather than within-trial sampling noise. As we show in the Supplementary Material (Figure S1B), however, such an assumption would not help, because increasing the variability in the drift rate parameter *d* with uncertainty across trials would have a similar effect on consistency and RT as increasing the within-trial diffusion noise (sampling variability) – higher variability will yield lower choice consistency and faster RT, contrary to the data (see Figure S1B in the Supplementary Material for a confirmation of this via simulations).

Another approach to the role of uncertainty on choice has been suggested in a recent study by Li and Ma (2020), who proposed a model in which the decision maker chooses based on a comparison of the subjective utility of each option. In their model, utility is a linear combination of the mean of the value estimate and the standard deviation of the value estimate (derived via Bayesian updating with the sequential evidence samples), suggesting that people explicitly prefer both options that are more valuable *and* those whose value estimates are more certain. Note that such a model would not modulate any of the standard DDM parameters as a function of value certainty, but would rather include a separate evidence accumulator variable that tracks the difference in certainty between choice options. We tested a model of this variety against our winning model and found that although it performed well on its own, it did not win the competition against the snDDM (see the Supplementary Material for the formulation of this Model 6 and the model comparison results). It would be interesting for future work to explore the similarities and differences between this type of model and the snDDM.

### Choice Confidence

While we have focused here on how value certainty affects choice consistency and RT, the empirical data also importantly show a marked and systematic effect of value certainty on choice confidence. In particular, higher certainty sum (C) and higher certainty difference (dC) both lead to higher choice confidence. This pattern raises a further challenge for most accumulation-to-bound style choice models that aim to account for both RT and choice confidence. For example, in the balance of evidence (BOE) type models (Vickers & Packer, 1982; De Martino et al, 2013), confidence corresponds to the difference in the activation of two accumulators that independently race to a response boundary. If we were to naively introduce option-specific value uncertainty in such models as additional processing noise, they would predict, contrary to the data, that the confidence becomes larger for options with more uncertainty (as the additional noise increases the average BOE). Similarly, if we were to model confidence using a DDM with collapsing boundaries (e.g., Tajima et al, 2016; Fudenberg et al, 2018), with confidence corresponding to the height of the boundary at the time the choice is made, naively introducing option-specific uncertainty would once again provide us with a prediction opposite from what we see in the data. For uncertain alternatives, there would be more noise in the evidence accumulation process, resulting in faster choices and therefore higher boundaries, and thus higher confidence (in fact, this would be true for any model that assumes that confidence decreases with RT; Kiani & Shadlen, 2009).

There are very few published value-based choice studies that simultaneously examined value certainty and choice confidence (but see Lee & Daunizeau, 2020, 2021; Lee & Coricelli, 2020; De Martino et al, 2013). We have not modeled choice confidence here, as there are many potential ways to do this, with substantial divergence among them (Vickers & Packer, 1982; Kiani & Shadlen, 2009; Pleskac & Busemeyer, 2010; De Martino et al, 2013; Moran, Teodorescu, & Usher, 2015; Calder-Travis, Bogacz, & Yeung, 2020; see Calder-Travis et al, 2020). Nevertheless, all of these models strive to predict a strong negative correlation between RT and choice confidence, as has been demonstrated in a plethora of experimental data. We note that in the data we examined, the impact of value certainty on choice confidence was essentially the reverse of its effect on RT (see Supplementary Material, Figure S7). While we did not explore this further, it suggests that a signal-to-noise DDM should also be able to capture the dependency of choice confidence on value certainty. Future work is needed to determine how signal detection style DDM variants might be extended towards an optimal unified account of choice consistency, RT, and confidence.

## Acknowledgments

We wish to thank Konstantinos Tsestsos and Giovanni Pezzulo for a critical reading of an early version of the manuscript and for helpful discussions, and Moshe Glickman for helpful discussion and advice on mixed model analysis. We thank Ian Krajbich for generously sharing his data with us. DL also wishes to thank Antonio Rangel for his generous support, encouragement, and helpful discussions. Finally, we wish to thank Jerome Busemeyer, for very helpful advice during the review process.

## SUPPLEMENTARY MATERIAL

### Simulation Results for the Impact of Value Certainty on Other DDM Parameters

In the Discussion section of the main manuscript, we explained why additional DDM variants (collapsing boundaries) or other assumptions about the impact of value certainty on the additional parameters in the standard DDM (drift rate variability, starting point variability) would not change the conclusions we presented here. Here we present simulation results that provide additional support to our conceptual explanations (Figure S1).

We start with the collapsing-boundary DDM variant. We show via simulations that if value uncertainty increased the sampling noise, this would decrease RT, just like in the fixed-boundary DDM (Figure S1A). We then consider the classic DDM with fixed boundaries and we examine whether value certainty could be introduced via an effect on either starting point variability or on drift rate variability (we consider the possibility that uncertainty might increase those variability parameters, so as to make to make the choice more random). We show via simulations that if value uncertainty increased the drift rate variability, this would decrease RT (Figure S1B). We also show via simulations that if value uncertainty increased the starting point variability, this would have no impact on mean RT (Figure S1C).

**Figure S1:**
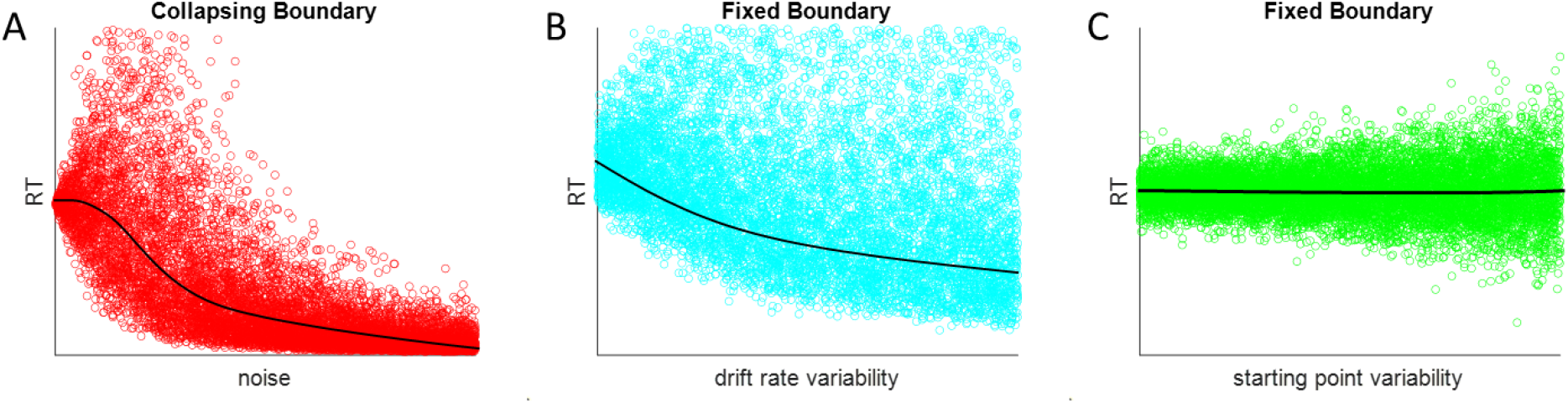
Simulated results demonstrating the relationship between additional DDM parameters and RT. For each plot, we simulated 10^4^ trials. Panel A show the relationships between diffusion noise and RT (see Figure 3, center panel), when the response boundaries collapse over time. Panel B shows the relationships between drift rate variability and RT. Panel C shows the relationship between starting point variability and RT. Each data point represents one simulated trial. The solid black lines indicate the mean RT as a function of the independent variable of interest.

### Response Time Distributions as a Function of snDDM Predicted Drift Rate

To verify that the snDDM can account for patterns in the full RT distributions (rather than only in mean RT), we performed an RT quantile analysis as advocated by previous studies (Ratcliff, 1979; Ratcliff, Van Zandt, & McKoon, 1999; Ratcliff & Tuerlinckx, 2002). Because we had many participants but only few trials per participant, we utilized a technique recommended by Ratcliff (1979), in which the independent variable of interest is summarized by its 10^th^, 30^th^, 50^th^, 70^th^, and 90^th^ percentiles for each participant and then averaged across participants. We present this summary for all subjects pooled together across studies in Figure S2, for the key snDDM variable of interest: (signal-to-noise difference) SN = V1*C1 – V2*C2. To obtain this data, we first separated the trials by a median split (within participant) and then divided the trials within each half of the median split by choice consistency (i.e., whether the higher-rated option was chosen or not). This provided us with four bins (high SN / inconsistent, low SN / inconsistent, low SN / consistent, high SN / consistent), within which we found the RT percentiles for each participant and then took the average across participants (Figure S2, left panel). The location on the x-axis represents the mean consistency level (across participants) for trials within each of the categories. To compare the empirical results with snDDM predictions, we simulated choice and RT using the real experimental input variables (V1, V2, C1, C2) and the best fit parameters for each participant. We then performed the identical analysis as with the experimental data (Figure S2, right panel). We found that the patterns across the full range of RT were similar between the simulated and the experimental data. In particular, in both, we can see the skew in the RT distribution (a larger separation for the higher RT quantiles compared the lower ones), and the fact that while the higher quantiles show a characteristic inverse U-shape, the lower quantiles appear to decrease monotonically with choice consistency.

**Figure S2:**
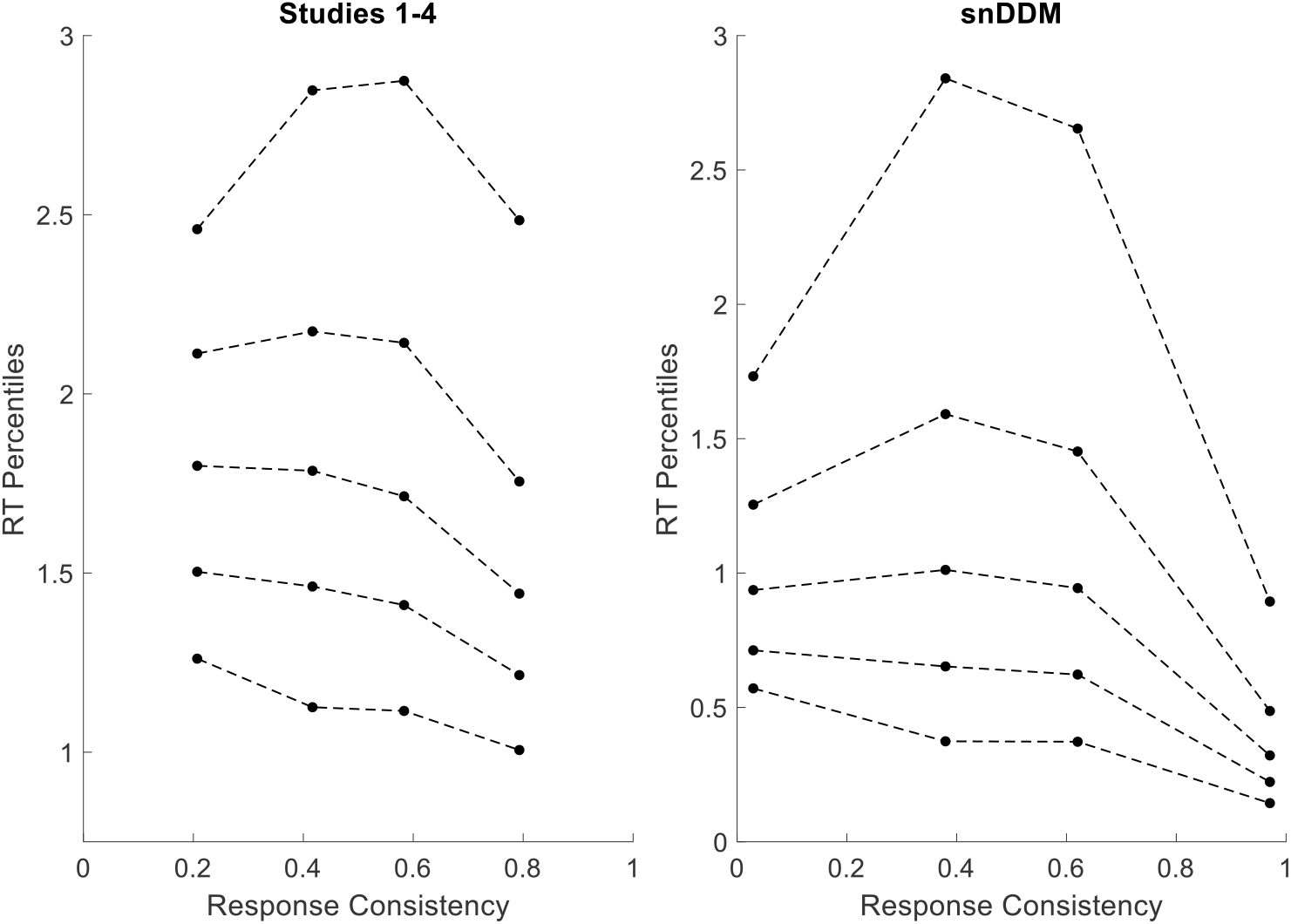
Quantile probability plots: RT percentiles as a function of response consistency (equivalent to choice accuracy), for different levels (median split) of the predicted signal-to-noise drift rate (SN). For each point, the x-coordinate represents the categorical bin {high SN / inconsistent choice, low SN / inconsistent choice, low SN / consistent choice, high SN consistent choice} and the y-coordinate represents the cross-participant mean of the RT quantile {0.1, 0.3, 0.5, 0.7, 0.9}. The left panel summarizes the experimental data (pooled across Studies 1-4) and the right panel summarizes the snDDM simulated output using real experimental input.

Below we illustrate the snDDM predicted relationship between C / dC and drift rate, which is inversely related to RT (Figure S3, left panel). The figure shows predictions across the entire (normalized) range of possible values of C1 and C2, of which C and dC are composed. We compare this to the experimental data (Figure S3, right panel), which summarizes RT (across participants) as a function of C1 and C2 (and thus C and dC). Note that while the match is not exact, the expected qualitative patterns hold between the theoretical and empirical data (i.e., RT is a decreasing function of both C and dC).

**Figure S3:**
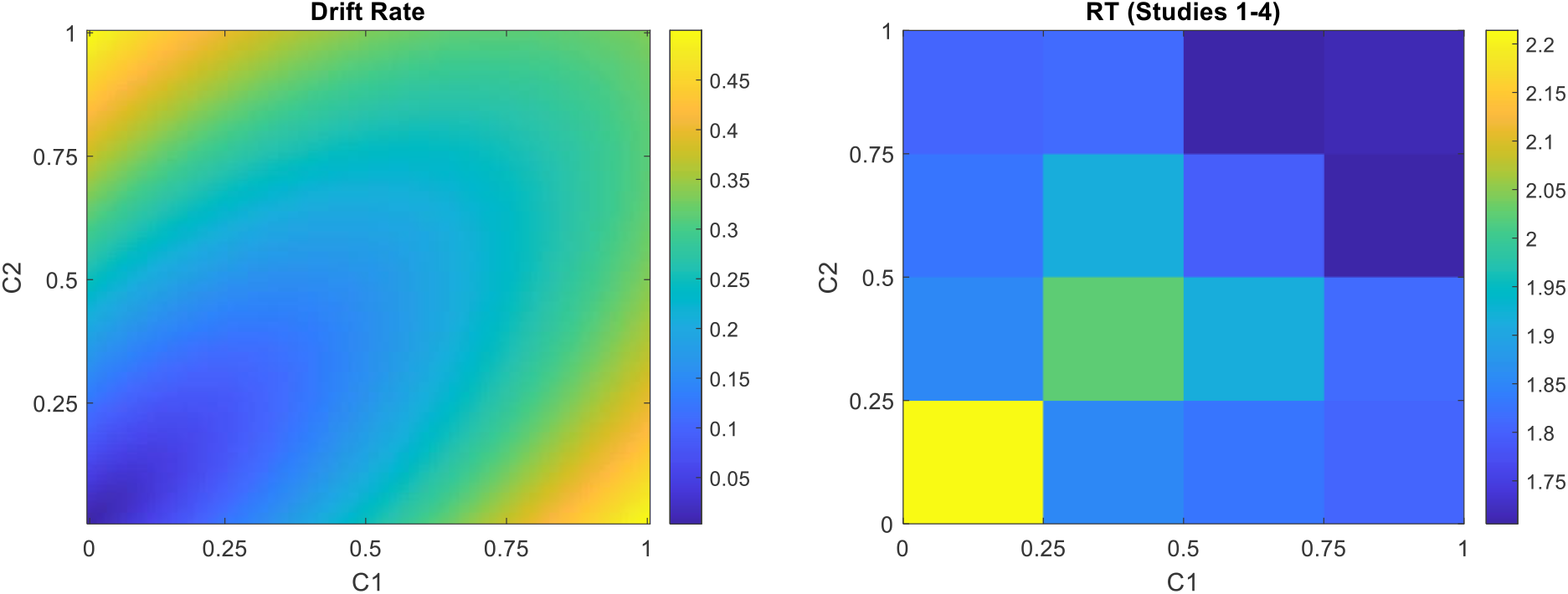
Drift rate under the snDDM is an increasing function of both dC and C (left panel). Averaged across the full normalized range of value (both V1 and V2), drift rate increased as either option increased in certainty in isolation (i.e., dC increased; moving upwards or rightwards in the plot), but also as both options increased simultaneously (i.e., C increased while dC remained constant; moving at a diagonal towards the upper right of the plot). Average RT (across participants) in Studies 1-4 shows the opposite pattern as the theoretical drift rate, as expected.

### Divergence from Ideal of the Experimental Input Data

In the main manuscript, we showed simulated results for how the different models we examined predict the relationships between certainty (sum and difference) and both choice consistency and RT using as input either theoretically ideal input data (Figure 4) or the actual empirical input data (Figure 5). The main differences between the ideal and actual data are that in the actual data, the variables did not cover the full available ranges, were not normally or uniformly distributed, and were partially correlated with each other. Specifically, the data was such that the distribution of C was highly rightward skewed, and the distribution of dC had the bulk of its mass close to zero (see Figure S4a). More importantly, value and certainty were not independent (see Figure S4b).

**Figure S4:**
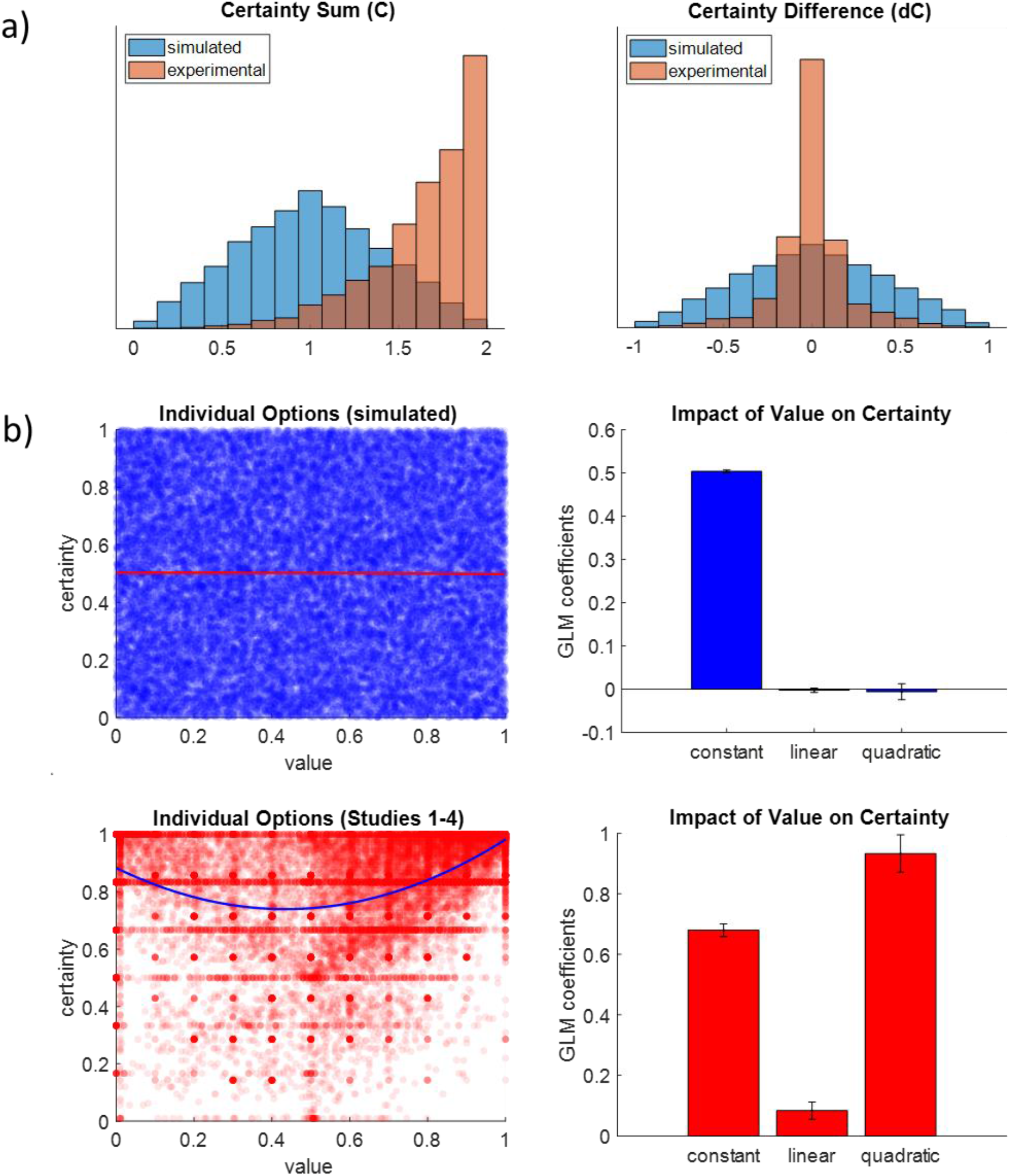
Divergence of experimental data from “ideal”. a) The distributions of C and dC obtained from the experimental data differed from those we used in our theoretical demonstrations of their impact on choice and RT under each of the models (Figure 4). b) The “ideal” data we used in our theoretical model simulations (in blue) evenly covered the entire value-certainty space, and were uncorrelated; the actual experimental data we used in the quantitative model comparison were concentrated in specific regions of the value-certainty space, and were significantly correlated.

### Impact of Overall Value on Response Time

Beyond the effects of value difference (dV), certainty sum (C), and certainty difference (dC) on RT, we also observed a strong effect of value sum (V). Specifically, higher V corresponds to lower RT (see Figure S5). As it turns out, the snDDM is able to predict this relationship (none of the other models can account for this; see Figure S5). This is especially interesting, given the fact that V itself never directly enters the model. Instead, the impact of V likely arises from the nature of the signal-to-noise structure of the model. The basic idea would be that the value signals for lower-valued options would be washed away by the detrimental impact of uncertainty, whereas the value signals for higher-valued options would be more resilient. For example, imagine two separate choice pairs: a) V1 = 0.2, V2 = 0.1, C1 = 0.8, C2 = 0.2; b) V1 = 0.9, V2 = 0.8, C1 = 0.8, C2 = 0.2. Under the standard DDM, both choice pairs would have the same mean evidence (e) dV = V1 - V2 = 0.1. Under the snDDM, the mean evidence for each pair is distinct: a) V1*C1 = 0.16, V2*C2 = 0.02 → e = 0.14; b) V1*C1 = 0.72, V2*C2 = 0.16 → e = 0.56. The larger mean evidence with larger overall value effectively makes the choice easier (i.e., the drift rate is higher), thus leading to lower RT. The same phenomena holds if the higher-valued option within each choice pair is the one with higher certainty, for example: a) V1 = 0.2, V2 = 0.1, C1 = 0.2, C2 = 0.8; b) V1 = 0.9, V2 = 0.8, C1 = 0.2, C2 = 0.8. Now, the mean evidence for each pair is: a) V1*C1 = 0.04, V2*C2 = 0.08 → e = -0.04; b) V1*C1 = 0.18, V2*C2 = 0.64 → e = -0.46. Again, the higher-valued choice is effectively easier (greater absolute evidence magnitude), and the choice can be made more quickly. Note that if both options within a choice pair had the same level of certainty, there would be no impact of V on RT, which explains why none of the other models we examined were able to account for that effect.

**Figure S5:**
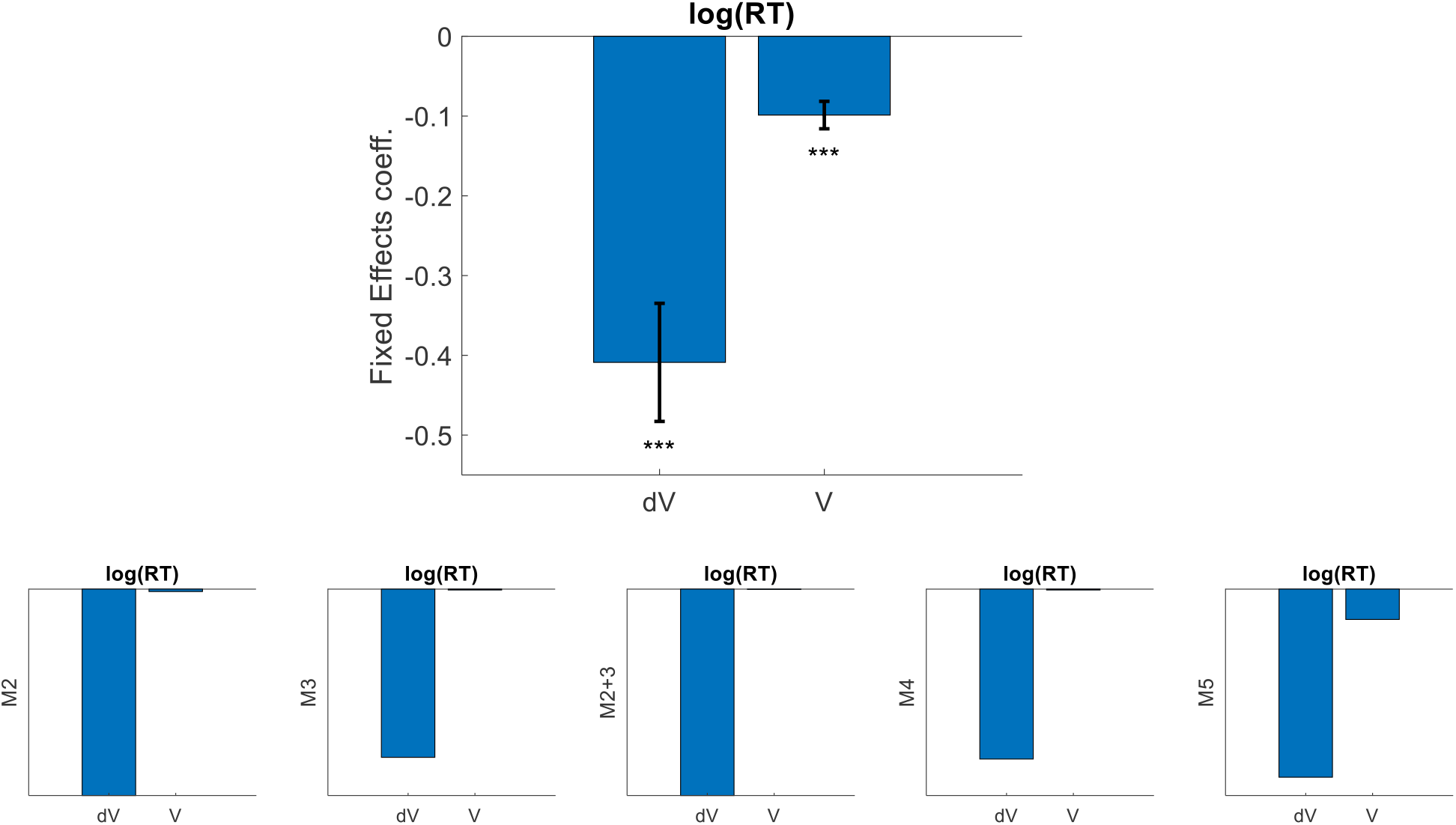
Impact of overall value on response time. The upper panel shows the fixed effects coefficients from mixed effect linear regression of value difference (dV) and value sum (V) on log(RT). Participants pooled across four studies, n=191; error bars represent standard errors; significance stars represent *** p < .001, one-sided t-tests. The bottom panel shows the same analysis for data simulated under each of the models we compared. Note that only Model 5 qualitatively matches the data.

Beyond the illustrative examples above, we also explored the relationship between V and RT in a more systematic manner. Specifically, we examined a 100x100x100x100 grid of V1, V2, C1, and C2 across a standardized range of 0:0.01:1. Within each cell in the grid, we calculated the drift rate as ashokV1*C1 – V2*C2ashok (we took the absolute value because it is the magnitude of the drift that impacts RT; the sign is irrelevant). We then averaged across all values of C1 and C2, leaving us with a two-dimensional array (V1 x V2) of calculated drift rates (see Figure S6). As expected, the drift rates were highest when ashokdVashok was largest (so when V1 and V2 had much different values). Beyond that, there was a clear positive relationship between drift rate and V, explaining the negative relationship between V and RT (which is inversely related to drift rate).

**Figure S6:**
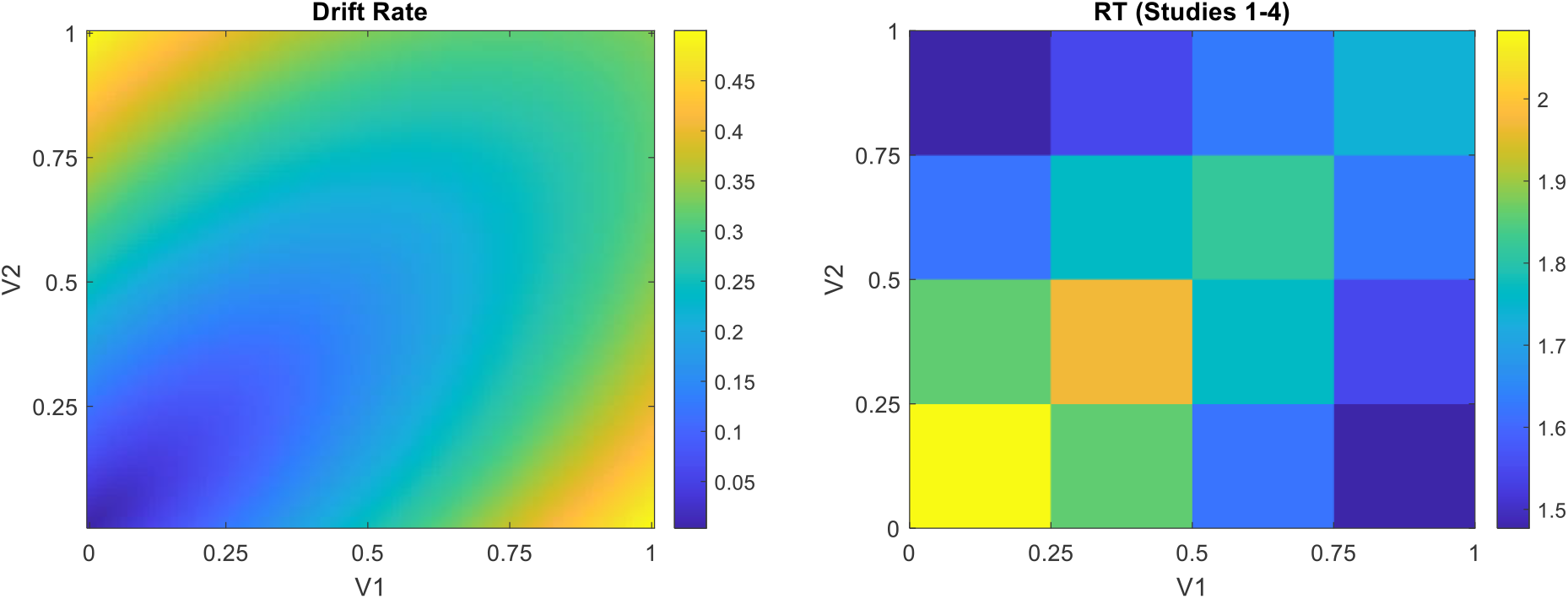
Drift rate under the snDDM is an increasing function of both dV and V (left panel). Averaged across the full normalized range of certainty (both C1 and C2), drift rate increased as either option increased in value in isolation (i.e., dV increased; moving upwards or rightwards in the plot), but also as both options increased simultaneously (i.e., V increased while dV remained constant; moving at a diagonal towards the upper right of the plot). Average RT (across participants) in Studies 1-4 shows the opposite pattern as the theoretical drift rate, as expected.

### Choice Confidence

In this study, we chose not to include choice confidence in our model predictions, as there is not currently an agreed-upon standard for doing so. Nevertheless, we did briefly examine this variable in those datasets that contained it (Studies 1-3; Lee & Daunizeau, 2020 2021; Lee & Coricelli, 2020). In general, choice confidence exhibited patterns qualitatively opposite to those exhibited by RT. Specifically, the mixed model fixed effects coefficients for dV, C, and dC were of similar magnitude as those for RT, but were all positive (whereas for RT, they were all negative; see Figure S7).

**Figure S7:**
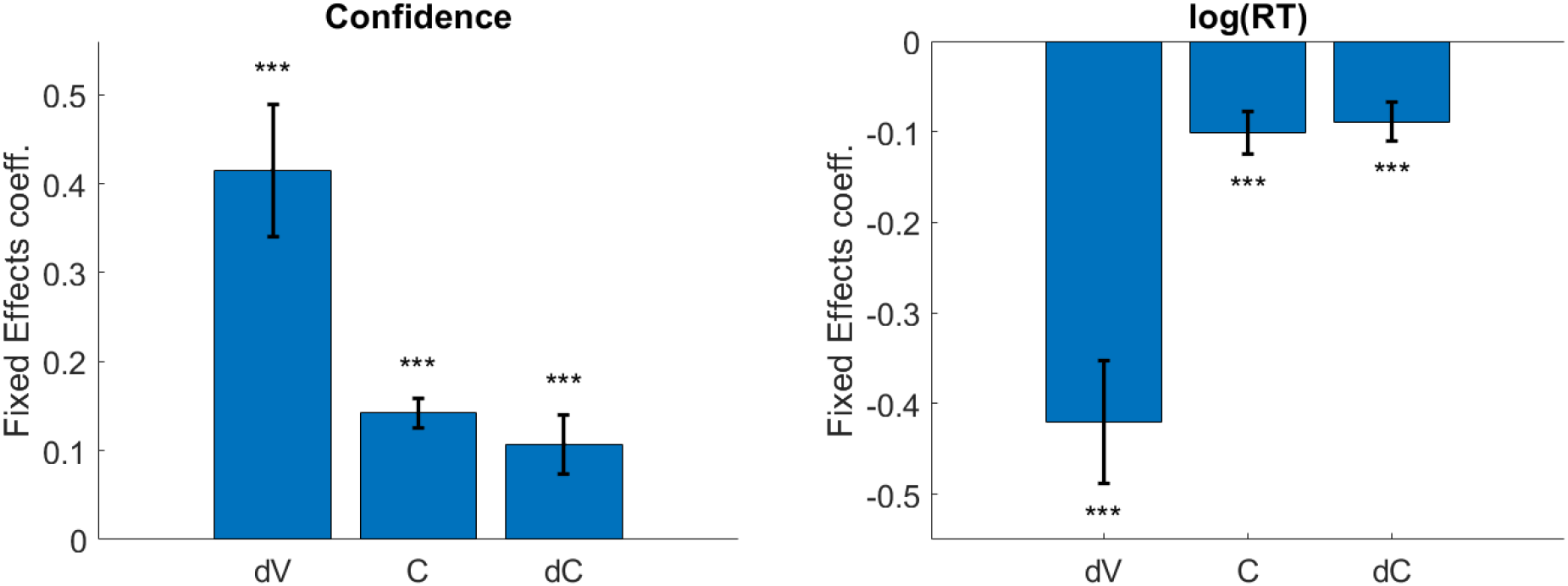
The fixed effects coefficients from mixed effect linear regression of dV, C, and dC on choice confidence (left panel) and log(RT) (right panel). Participants pooled across three studies, n=155; error bars represent standard errors; significance stars represent *** p < .001, one-sided t-tests.

### DDM Median Split Parameter Comparison

To provide additional corroborating evidence that value certainty should relate to the drift rate and not the diffusion noise parameter, we performed classic median-split analyses on both the sum and the difference of the value certainties of the choice pair options. For each participant across all studies, we divided the trials based on a participant-specific median split of either certainty sum or certainty difference. We then fit the classic DDM separately for each of the subsets of data and compared the fitted parameters across subsets. As shown in Figure S8, in both the certainty sum median split and the certainty difference median split, the drift rate parameter was larger in the high versus low subsets; the noise parameters did not differ across subsets. This shows that our conclusion that Model 5 (with drift rate adjusted by certainty) is valid, while Model 2 (with diffusion noise adjusted by certainty) is not.

**Figure S8:**
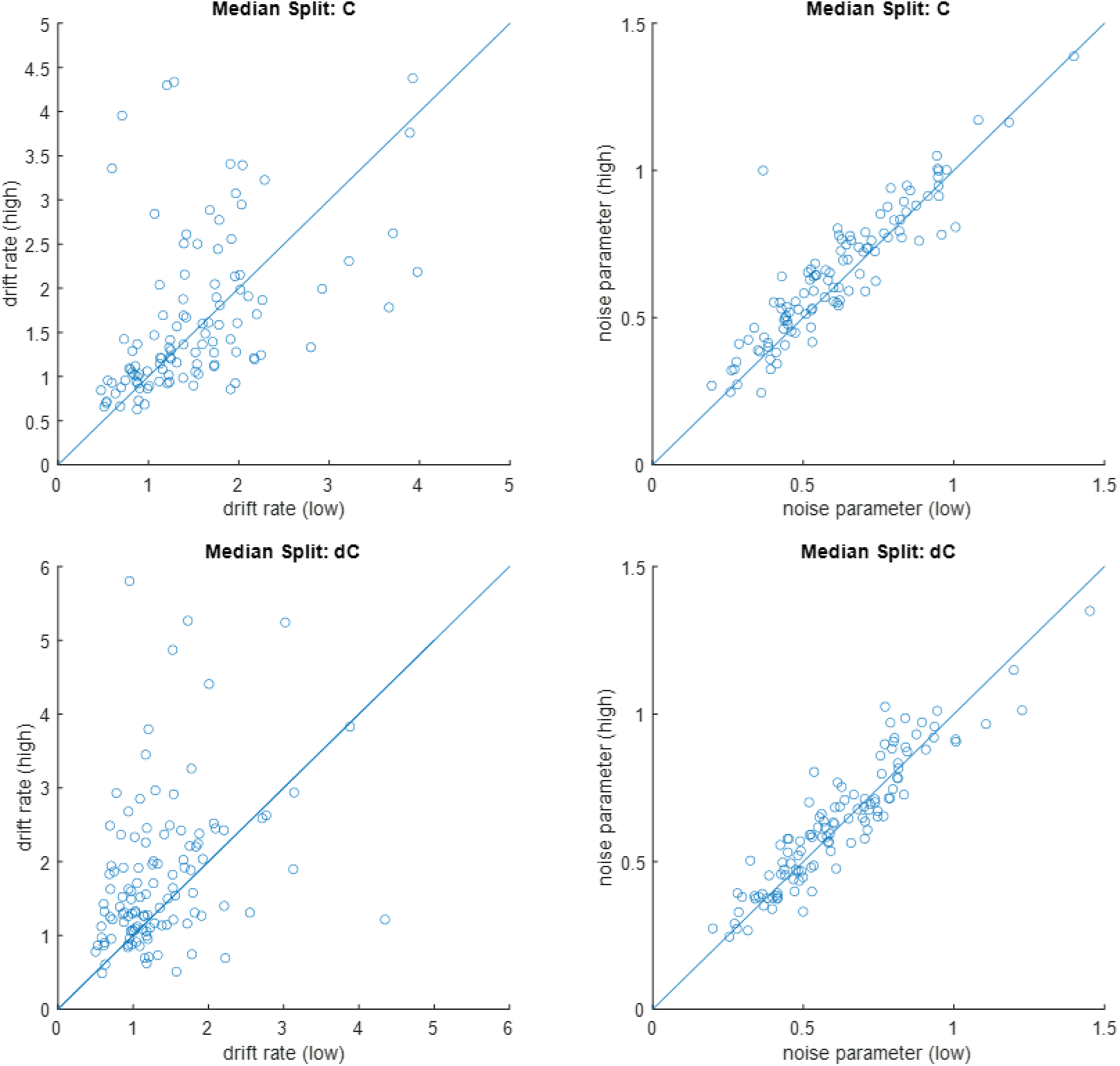
DDM parameter comparison across median splits of certainty sum (top panels) and certainty difference (bottom panels). Each data point represents one participant. Across the population, the fitted drift rate parameters were larger for the high certainty trials as well as for the high certainty difference trials (left panels); the fitted diffusion noise parameters were not different across high and low certainty or certainty difference (right panels).

### Starting Point Bias towards Left or Right Option

As a simple check to see if the participants in the studies we examined had a tendency to favor the options presented on the left or right side of the screen (independent of option values), we calculated a simple percentage of the total choice trials for which the participants chose the left option (since the value of each option was orthogonal to its physical location on the screen). Overall, the participants chose the left option on 50% of the trials. At the individual level, 91% of participants chose the left option on 40-60% of trials. We tested that these individual deviations from 50% were due to chance rather than bias. We thus calculated, for each participant, the standard deviation of a binomial distribution 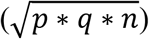 with a 50% probability (p = q = 0.5) and n = number of trials for that participant (which varied across datasets). We then computed a t-statistic for each participant by subtracting n/2 from the number of left responses and dividing by the standard deviation. Using t = +/− 1.96 as a cutoff for a 95% confidence interval that the leftward response rate for each participant was not statistically different than 0.5, we found that only 8% of participants fell outside of this confidence interval. We thus concluded that there was no meaningful side response bias in our data.

### Model 6: Separate Accumulator for Certainty

In this study, we examined a number of alternative ways in which the traditional DDM parameters (drift rate, diffusion noise, and response boundary) could be modulated by option-specific value certainty. Yet a fundamentally different way in which the DDM could include value certainty is in the form of an independent evidence accumulator—a secondary drift, with a rate proportional to the difference in certainty between the choice options. In this way, the decision about which option to choose would be influenced both by which value estimate was higher (via the primary, standard drift) and by which value estimate was more certain (via the secondary, novel drift). The secondary drift might represent an aversion to uncertainty, where the deliberation process would be both slowed by the uncertainty and the decision pulled towards the more certain option. This would imply that the decision maker prefers options that are more valuable, but also options for which the value is more certain. Other ongoing research has proposed a model based on a similar principle (Li & Ma, 2020). Under our version of this framework, the equation governing the DDM process would be:

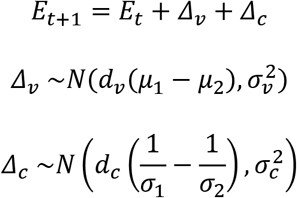

where Δ_v_ and Δ_c_ are the incremental evidence for value and certainty, respectively, and d_v_ and d_c_ are scalars (it is assumed that d_v_ will always be positive, but d_c_ could take either sign). The inclusion of a secondary drift rate (in essence, a parallel evidence stream that monitors value certainty rather than value itself) can have either a positive or a negative impact on both choice and RT, depending on whether the option with the higher mean evidence stream is the same as or different than the option with the higher evidence reliability (as well as on the sign of d_c_). This can be seen by examining the (revised) DDM equations for choice probability and expected response time:

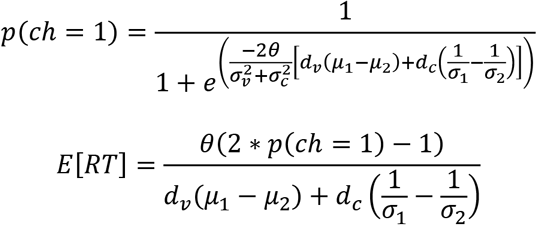

Here, the secondary drift rate will result in more consistent and faster choices if the sign of μ_1_-μ_2_ is the same as that of 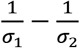, but it will result in less consistent and slower choices otherwise. This, of course, is under the assumption that the decision maker is uncertainty averse (i.e., d_c_>0). If the decision maker were uncertainty seeking (i.e., d_c_<0), the opposite predictions would hold.

Model 6 could account for the empirical relationship between certainty difference (dC) and both choice consistency and RT. However, Model 6 would fail to account for the dependency of consistency and RT on certainty sum (C) (see Figure S9).

**Figure S9:**
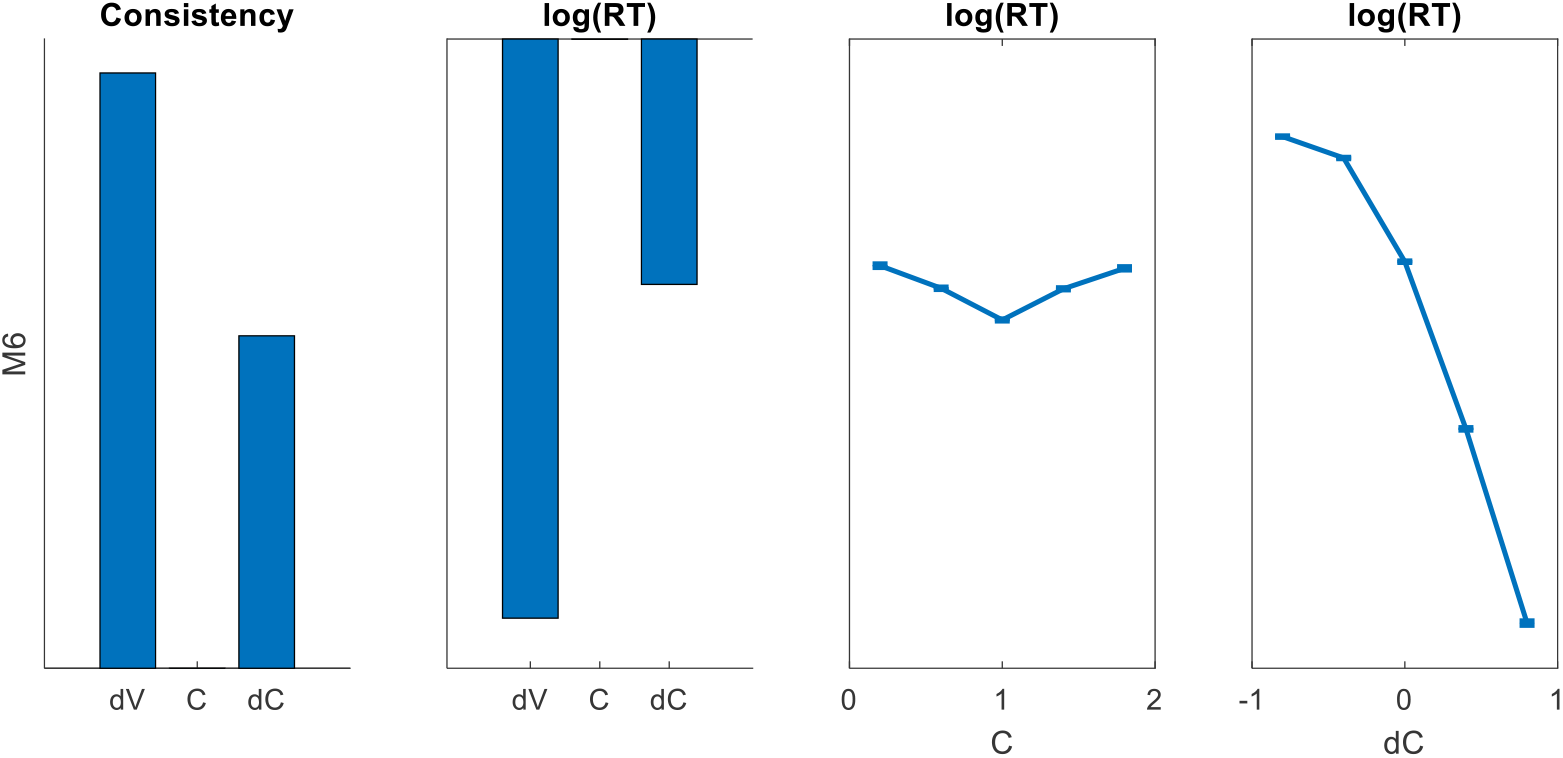
Qualitative predictions of the effects of value difference (dV = V1 - V2), value certainty (C = C1 + C2), and certainty difference (dC = C1 - C2) on choice consistency and log(RT) (shown for the best fit parameters; bar heights represent mean GLM beta weights based on 10^4^ simulated trials for Model 6). The log(RT) data were z-scored prior to binning for the quantile curve plots.

To test the ability of this alternative formulation to explain the data, we ran a formal model comparison between our winning model (Model 5) and Model 6. Model 5 dominated the model comparison (exceedance probability = 1), with Model 6 receiving a trivial amount of support (approximately 8% of participants pooled across all datasets; see Figure S10).

**Figure S10:**
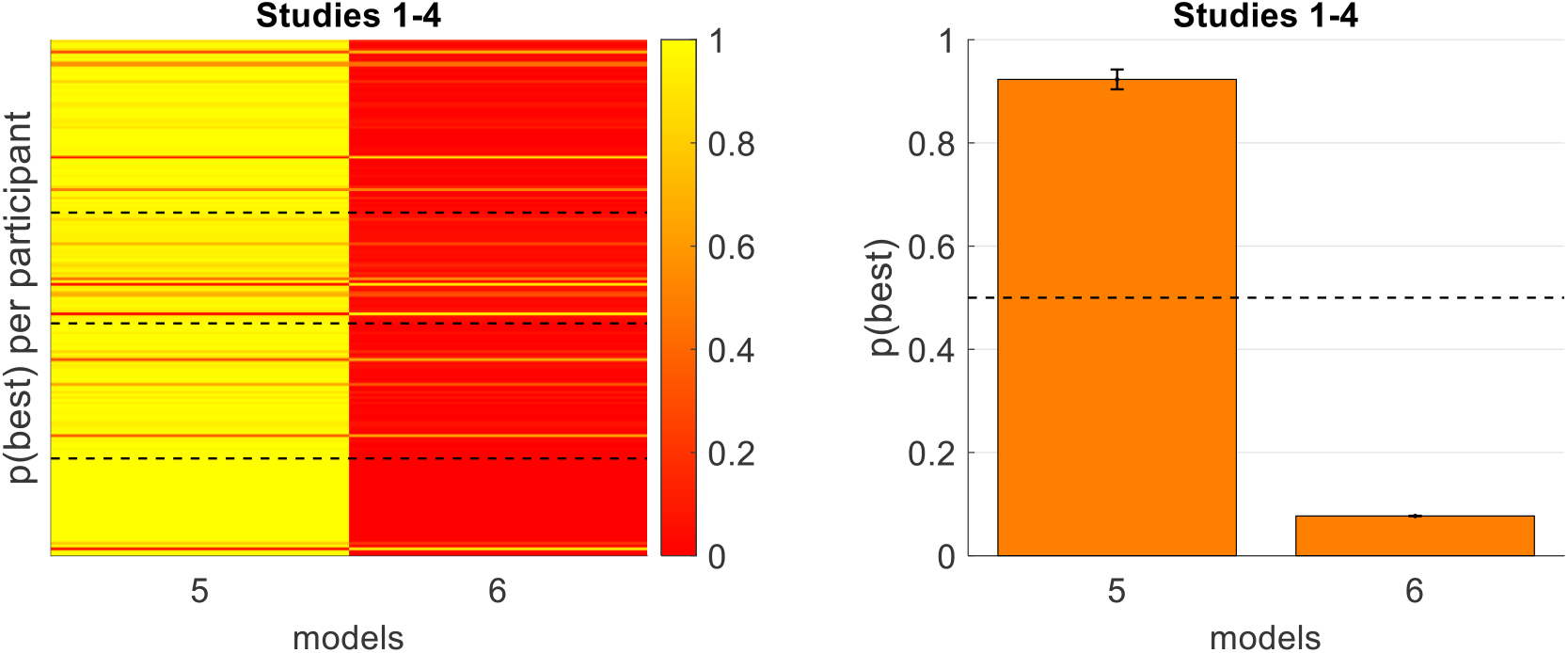
Model comparison results for Model 5 vs Model 6 (same format as Figure 6).

### Best-Fitting Parameters

Across all models, we observed that the best-fitting parameters were fairly consistent (see Figure S11), although there was some variation in the distributions (as expected). The individual parameters were closely correlated within participant across models.

**Figure S11:**
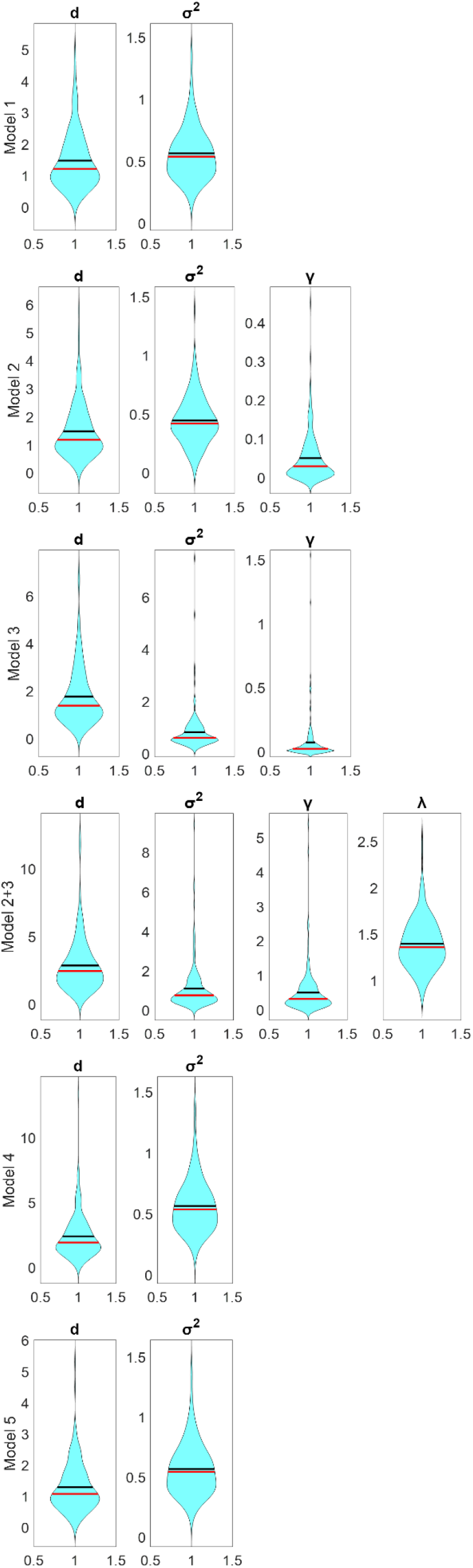
Distributions of best-fitting model parameters, across all participants. Violin plots represent cross-participant distributions of GLM beta weights; black lines represent cross-participant mean values, red lines represent cross-participant median values.

We also checked to see if the key DDM parameters, drift rate *d* and diffusion noise σ^2^, were correlated. In particular, we examined the basic DDM (which has already been validated numerous times; our Model 1) and our winning model (Model 5). As shown in Figure S12, there is a positive correlation between the two parameters, both in Model 1 and in Model 5.

**Figure S12:**
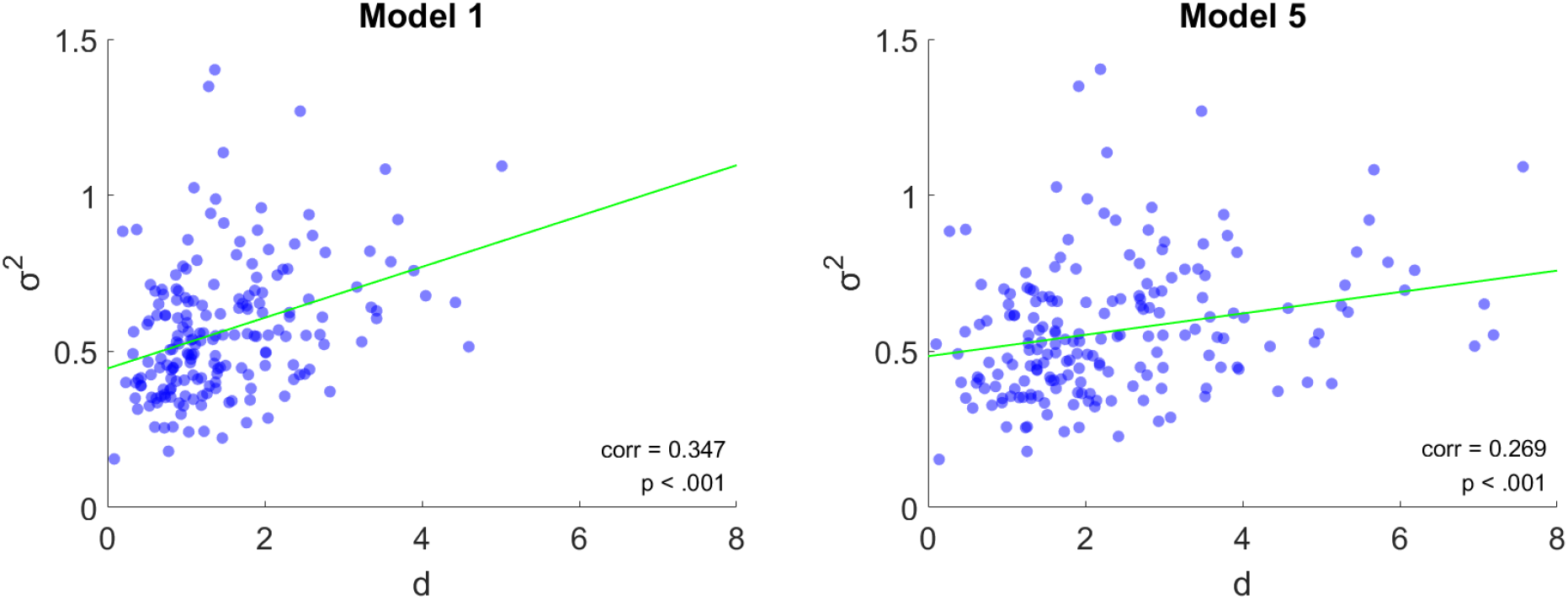
There was a positive relationship (across participants) between the best-fitting drift rate d and diffusion noise σ^2^ in both Model 1 (left panel) and Model 5 (right panel).

## Footnotes

1 According to Tajima et al (2016) and Fudenberg et al (2018), the optimal policy requires a collapsing rather than fixed boundary. The collapse of the boundary results, however, from auxiliary assumptions about the cost of deliberation time to the decision maker. Since these assumptions can be debated, and since there is an intensive debate on whether experimental data best support DDM with fixed or collapsing boundaries (Evans, Trueblood, & Holmes, 2020; Hawkins et al, 2015; Palestro et al, 2018), we consider both cases in this study. We will start with the simple fixed boundary DDM, which is the most prominent version in the literature, and we will then show that our conclusions are unchanged for the collapsing boundary variant (see Figure S1 in the Supplementary Material).

2 As shown in Figure 2, we regressed the various measures of certainty on log(RT) rather than on RT. This is because whereas the RT variable is not normally distributed (it has a relatively long right tail), the transformed log(RT) variable is normally distributed and is thus better suited for statistical testing (see Glockner & Betch, 2008). Our conclusions hold if we use RT (without the log-transformation).

3 In another prominent evidence accumulation choice model – the *linear ballistic accumulator* (LBA) model (Brown & Heathcote, 2006) – the within-trial noise is replaced by between-trial noise in the starting point and drift rate (in the DDM, such between-trial variability is sometimes introduced in addition to the within-trial noise; Ractcliff & McKoon, 2008). Increasing those components with option-uncertainty, however, would have a similar effect as increasing the within-trial diffusion noise in the DDM (i.e., faster responses and reduced accuracy; see the Supplementary Material).

4 In the standard version of the DDM, the RT distribution for correct and incorrect responses is identical. In a more complex version, additional variability parameters are introduced that allow it to account for asymmetries between the RT distributions of correct and incorrect responses (see Ratcliff & McKoon, 2008 for a review). We only consider the standard DDM without these variability parameters, as they do not change the impact of value certainty on consistency and RT illustrated in Figures 2 and 3.

5 Because this system of equations is over-parameterized, one of the free parameters must be fixed for a practical application of the equations. In this work, for simplicity, we fix to a value of 1 (when fitting the models) the boundary parameter θ.

6 It is important to clarify that we do not see Model 2+3 as an actual application of DFT to the task of deciding between food items, since DFT includes additional processes. For example, DFT incorporates approach/avoidance, which corresponds to a “leak” parameter in the value integration (this transforms the diffusion from a Wiener process to an Ornstein-Uhlenbeck process, which requires more complex fitting procedures). Additionally, under DFT, the diffusion noise also depends on the correlation between the value fluctuation of the alternatives — a measure that we do not know how to assess in the experimental paradigm that we examine in this study. Model 2+3 is thus only an exploration of one specific DFT-inspired principle, within the framework of simple DDM variants. We also note that when fitting this model, we do not fix θ to 1 as we do in the other models. Here, the response boundary is dependent on the diffusion noise, so it is helpful to leave θ as a free parameter in order not to over-constrain the model.

7 Specifically, multiplying the drift rate by *x* is mathematically identical to dividing the boundary by *x* while simultaneously dividing the noise by *x^2^*.

8 In the actual experimental data, value and certainty ratings are correlated, and neither are uniformly distributed (see Figure S4 in the Supplementary Material). Thus, these predictions differ somewhat from what the models predict based on the actual input data (see Figure 5 below). Nevertheless, we show the theoretical predictions in Figure 4 to better illustrate the differences between the model variants.

9 Under Model 5*, most participants showed a concave impact of value certainty on evidence accumulation, with 70% of participants having α < 1 (28% had α > 1). Across participants, the median recovered α was 0.68 with a median absolute deviation of 0.46. This suggests that the marginal impact of value certainty is greater for options with relatively low certainty compared to those with relatively high certainty, for most participants. However, we note that these results should be interpreted with caution, because the parameter estimates of the α parameter were not very reliable. The parameters for each participant were estimated using very few trials, which presents a challenge to accurately fitting exponential parameters in addition to the linear ones. In particular, our parameter recovery exercise showed that the exponent parameters were often overestimated.

10 The empirical data exhibits few clear patterns: consistency increases with both C and dC, RT decreases with both C and dC; these relationships are mostly monotonic, although there is some noise in the lower extreme of the dC range with respect to consistency. All models appear capable of qualitatively matching the C-consistency relationship. Model 5 is capable of qualitatively matching the dC-consistency relationship, while the other models predict a flatter relationship with a dip when dC is close to 0. All models other than Model 2 appear capable of qualitatively matching the C-RT relationship. Model 5 is capable of qualitatively matching the dC-RT relationship, while the other models predict flat or non-monotonic relationships.

11 From this point forward, we will refer to the best performing model, Model 5, as the signal-to-noise DDM, or *snDDM*.

12 The DFT model can account for faster RT with higher values, as a result of its approach-avoidance assumption that modulates the integration leak parameter; with higher values, this is an approach situation that reduces the diffusion leak, resulting is faster choices (Busemeyer & Townsend, 1993; see Table 12).

